# Diurnal rhythms of choice: a novel state-dependent drift diffusion model uncovers time-dependent changes in rat decision making

**DOI:** 10.64898/2026.05.25.727672

**Authors:** Ryan A. Senne, Hongjie Xia, Helene F. Duebel, Quan Do, Gary A. Kane, James Fourie, Steve Ramirez, Benjamin B. Scott, Brian D. DePasquale

## Abstract

2

Time-of-day severely impacts human decision-making, with real-world consequences. Studying shifts in decision-making strategy requires controlled, long timescale behavioral measurement and analyses that can extract insight from time-varying behavior. We introduce two complementary advances to address this gap: an autonomous 24-hour training facility for continuous behavioral measurement during decision-making and an interpretable modeling framework that captures non-stationary decision dynamics from reaction times and choices. Rats were trained on a visual evidence accumulation task across months, generating over a half million trials spanning the circadian period. Our model revealed latent behavioral states characterized by distinct evidence accumulation parameters, including differences in drift rate, bias, and decision-commitment time. These states recur across days and align with feeding schedules and the light–dark cycle, producing periodic fluctuations in performance over 24 hours. Together, these results demonstrate how continuous behavioral sampling combined with generative modeling uncovers long-timescale structure in decision-making obscured by stationary analyses.

**HIGHLIGHTS:**

- 24-hour live-in operant system allows autonomous training in cognitive tasks across months
- 24-hour measurements reveal that rat performance fluctuates with time of day
- Novel DDM-HMM framework identifies reaction time and accuracy shifts across multiple timescales
- DDM-HMM captures serial dependence in decisions that classic models ignore

## 4 INTRODUCTION

Human and animal decisions vary systematically with behavioral context. Over the course of a day, internal states such as arousal, motivation, and fatigue fluctuate alongside changing environmental demands, producing time-dependent shifts in decision strategy. These fluctuations are not merely incidental: in real-world settings where decisions must be made continuously, state-dependent variation can have substantial behavioral consequences. For example, sleep-burdened doctors commit more medical errors and drowsy driving leads to more accidents.^1–3^

Studying how decisions vary across time requires controlled experiments spanning extended time-scales and analysis approaches that can identify temporal changes in decision strategy. Standard approaches cannot meet these demands. Many studies of decision-making in rodents are conducted within short, fixed daily sessions, typically lasting one to two hours.^4–8^ While efficient, this approach measures behavior from a restricted temporal window that obscures slower fluctuations in behavior.^9^ This constraint has important consequences, as circadian and sleep–wake homeostatic processes modulate vigilance, attention, arousal, learning, and performance.^10–16^ Consequently, apparent differences in decision-making across animals, tasks, or species may reflect the environmental conditions under which behavior is measured—such as time of day or motivational state—rather than intrinsic differences in cognitive or perceptual processing. To mitigate these concerns, some laboratories employ reversed light–dark cycles so that behavioral testing coincides with the animals’ subjective night, when rodents are typically most active.^10,17,18^ However, this approach introduces complications, including prolonged entrainment periods and potential disruptions to naturalistic behavioral rhythms.^10^

To address these limitations, we developed an autonomous, 24-hour living-and-training system that allows rats to perform a perceptual decision-making task continuously. This approach enables behavioral measurements spanning the full circadian cycle under stable experimental conditions. Using this system, we collected a large dataset of self-paced choices and reaction times from rats performing a visual perceptual integration task across complete 24-hour cycles.

The scale and continuity of these data revealed substantial variability in behavior over time, motivating the need for analytical tools that can provide insight into the underlying causes of this variation. Classical evidence accumulation models, such as the drift diffusion model, provide frameworks for interpreting choices and reaction times but assume that behavior is generated by a time-invariant process.^19,20^ Recently-developed state-based models capture slowly-evolving changes in behavior over longer timescales but often ignore reaction time information that yields critical insight into the mechanistic underpinnings of the decision process.^21–24^ To bridge this gap, we developed a statistical framework that combines the advantages of state-dependent models and the drift diffusion model, allowing for latent changes in decision state while retaining mechanistic interpretability. This approach provides a principled framework for examining how perceptual decision-making strategies vary over long timescales and across internal and external states.

We applied our framework to our unique dataset and conventional free-response datasets and found behavioral performance and its computational interpretation are strongly shaped by an animal’s circadian and motivational state, providing critical insight into decision-making behavior during a naturalistic setting.

## 5 RESULTS

### 5.1 Implementation of a live-in operant system for cognitive assessment

To enable continuous assessment of behavior across the circadian cycle, we developed RatAcad, a live-in operant training facility that allowed rats to engage freely with a cognitive task over a full 24-hour period under constant task parameters (fixed light–dark cycle and feeding schedule) with uninterrupted sucrose-based fluid reward (**Figure 1A**). This system was inspired from previous large-scale behavioral apparatuses, but notably diverges in its ability to *jointly* house and train animals.^25^ Behavioral data were acquired continuously and synchronized daily, while human interaction was limited to essential husbandry to minimize experimenter-induced effects.

**Figure 1:**
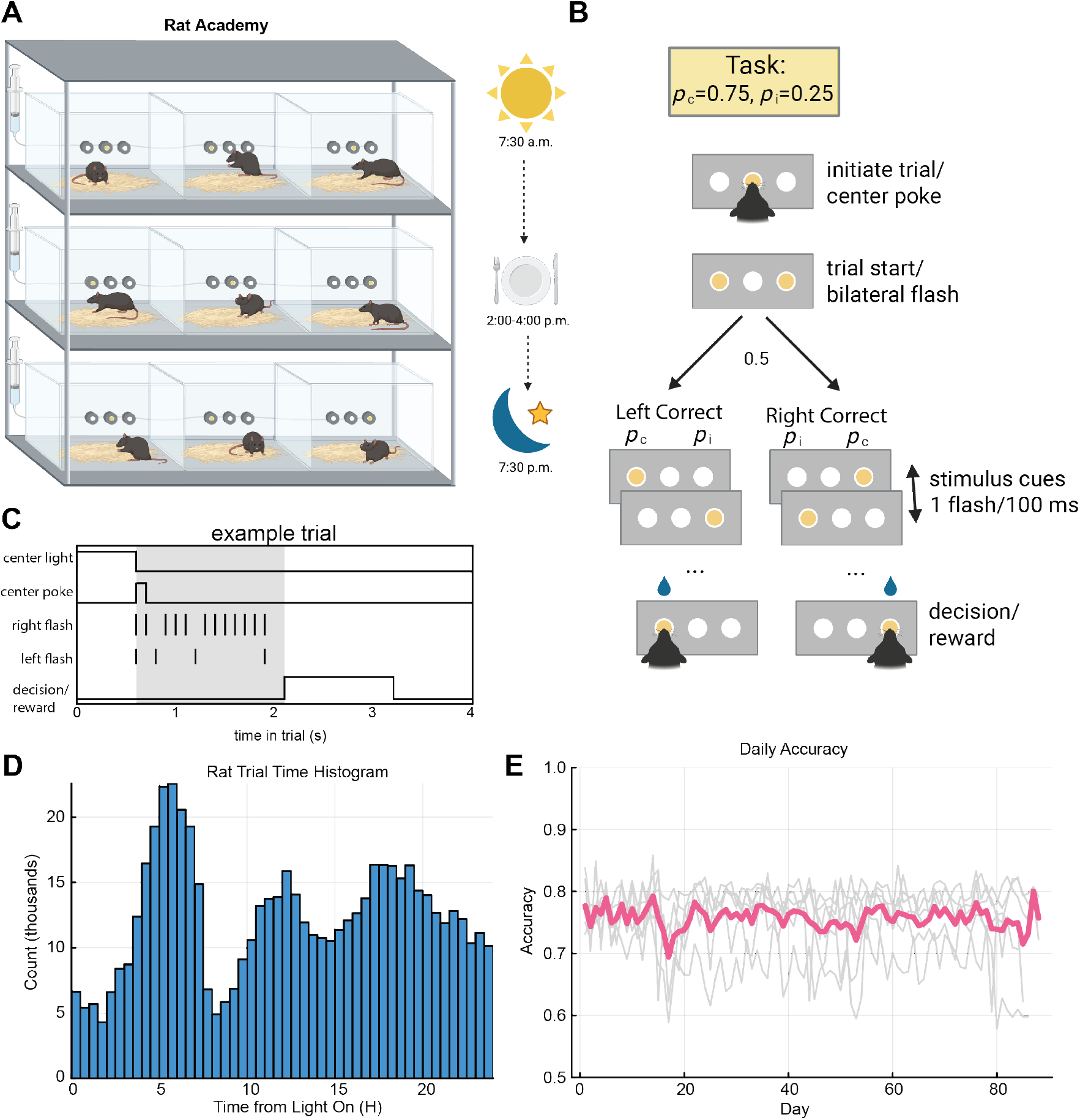
Automated evidence accumulation task and chamber. **(A)** Diagram of the final stage of the visual evidence accumulation task. The correct flash port (*p*_*c*_ = 0.75) is selected with equal probability between the left and right decision ports on each trial following trial initiation. Rats can observe stimulus flashes (up to 8 s total) at will and report their choice voluntarily. **(B)** Schematic of an example trial. **(C)** 24-hour operant chamber. Rats live in and continuously perform the visual evidence accumulation task within the same chamber. Lights are on from 7:30 AM to 7:30 PM, and rats are fed daily between 2:00 and 4:00 PM. **(D)** Trial-time histogram showing the number of trials collected in 30-minute bins across all rats over the 24-hour cycle. **(E)** Daily average accuracy.

To understand how animal decision-making performance varies across the 24-hour day, we trained rats on a visual-based perceptual integration task.^26^ In this free-response paradigm, animals performed a two-alternative forced-choice task in a three-port operant chamber. Each trial began with an initiating center poke, after which animals were free to choose the left or right port at any time during a cue period. During this period, the two side lights flickered stochastically with unequal probabilities (75% vs. 25%), and rats were rewarded for choosing the side associated with the higher probability of flashes (**Figure 1B–C**).

Before inclusion in the present analyses, animals were trained using a structured six-stage behavioral protocol to ensure stable task engagement and performance (**Supplemental Figure 1A**). Using this protocol, we trained 18 rats (12 male, 6 female) yielding 562,645 behavioral trials (**Supplemental Figure 1B**). This data was collected over long time periods, spanning weeks to months, which would have been comparatively difficult to collect using standard training paradigms. We collected thousands of trials at all hours of the day (**Figure 1D**); animal performance remained stable across days (**Figure 1E**). Together, these improvements allow for more rapid data collection, reduced need for human interaction, and the observation of changes across longer timescales e.g., days and months. Perhaps more importantly, it allowed us to examine how decision-making changed across these varying timescales.

### 5.2 The 24-hour cycle robustly modulates accuracy, reaction time, and engagement

We observed task-related behavioral fluctuations across the 24-hour period (**Figure 2A–C**). To quantify time-of-day effects, we fit generalized additive mixed models (GAMMs) using time of day as a covariate for each behavioral variable (accuracy, RT, and trial rate). The fixed effects of these models (representing population-level trends) revealed systematic modulation by diurnal phase (**Figure 2D–F**). Accuracy varied significantly across the 24-hour cycle (edf = 16.7, *χ*^2^ = 859.21, *p <* 10^−16^). Reaction times showed a complementary pattern (edf = 13.2, *F* = 41.55, *p <* 10^−16^), with fastest responses during feeding and slowest during the dark phase. Trial rates also exhibited strong temporal modulation (edf = 13.0, *F* = 45.20, *p <* 10^−16^), with animals performing fewer trials during feeding and showing maximal engagement in the dark period. These results demonstrate that behavioral performance, response speed, and task engagement are all tightly coupled to the local environmental conditions.

**Figure 2:**
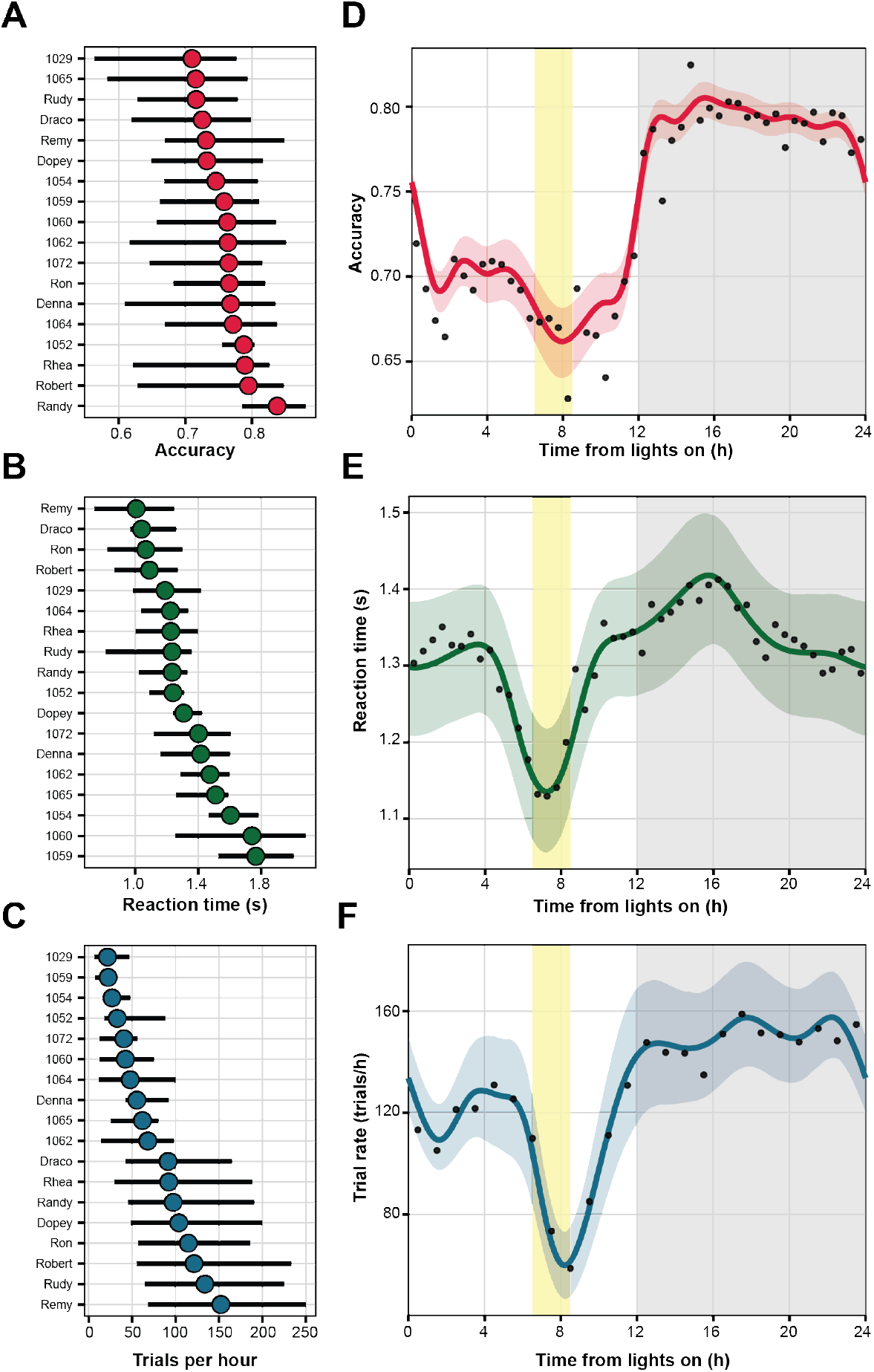
Individual variability and circadian modulation of behavior. **(A–C)** Dot-and-whisker plots showing variability across animals for **(A)** accuracy, **(B)** reaction time (RT), and **(C)** trials per hour. Dots indicate the mean for each animal, and horizontal lines indicate the range. **(D–F)** Fixed effects from generalized additive mixed models (GAMMs) illustrating the time course of behavioral performance across the 24-hour cycle. **(D)** Accuracy varies significantly with time, reaching a minimum during feeding (yellow shaded region) and a maximum during the dark phase (gray region). **(E)** Reaction times show a similar pattern, being shortest near feeding and during the dark phase. **(F)** Trial rates also exhibit circadian modulation, with engagement dropping during feeding and peaking during the dark phase.

Inspection of subject-specific GAMM fits revealed inter-animal variability in the magnitude and temporal dynamics of the effect across all three metrics despite a consistent qualitative structure (**Supplemental Figures 2–4**). Most animals exhibited a transient modulation around the feeding window, followed by a gradual recovery or overshoot, but the depth of the trough, latency to peak, and post-feeding trajectory varied between individuals. Notably, some animals showed sustained post-feeding changes whereas others returned rapidly to baseline, indicating heterogeneity in both sensitivity and recovery dynamics. These individual differences were evident even when overall exposure and trial counts were comparable, suggesting that the population-level effects are not driven by a small subset of animals but instead reflect a common pattern expressed with animal-specific timing and amplitude.

### 5.3 The DDM-HMM: a novel state-space model for serially-dependent evidence accumulation

The 24-hour modulation of accuracy, reaction time, and trial rate violates a core assumption of many standard sequential sampling models: that all trials are generated from a single, time-invariant process. To address this we developed a novel state-space model, the Drift Diffusion Model–Hidden Markov Model (DDM-HMM), to capture structured, trial-to-trial variability in decision-making while retaining the interpretability of classical evidence accumulation models. The DDM-HMM links reaction time distributions and choice outcomes to latent cognitive parameters, including drift rate, decision boundary, and starting point, while loosening the assumption of time invariance imposed by traditional models like the DDM.^19,20,27^ The DDM-HMM assumes reaction times and choices are serially correlated and generated by stochastic transitions among a discrete set of *K* latent DDMs. On each trial, a reaction time–choice pair is distributed according to a state-dependent Wiener First Passage Time (WFPT) distribution,

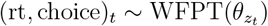

where each latent state *z*_*t*_ indexes a distinct set of DDM parameters *θ*. For each latent state *k*, the parameter vector is *θ*_*k*_ = {*B*_*k*_, *v*_*k*_, *a*_0,*k*_, *τ*_*k*_} where *B* is the bound height that determines when a decision is committed to, *v* the drift rate that defines the strength of accumulated evidence, *a*_0_ the starting-point bias that describes choice bias that is stimulus-independent, and *τ* the non-decision time, which defines the time between decision commitment and decision report. Recent studies determined that decisions by mice performing a similar decision-making task reflected abrupt temporal transitions between a small number of discrete decision-making strategies.^24^ Following this insight, we model the temporal evolution of the latent state according to a first-order Markov process:

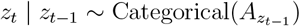

The model segments behavior into discrete, temporally-dependent decision states, each corresponding to a distinct DDM. (See methods for details).

A key innovation of our approach is the use of reaction time information in a state-dependent decision-making model. Choice-only state-switching approaches, such as the generalized linear model hidden Markov model (GLM-HMM) described above, partially address non-stationarity by segmenting behavior into discrete epochs, but they do not incorporate reaction time information during inference when reaction times are germane to the task.^23,24^ By leveraging the full first-passage-time distribution, the DDM-HMM disambiguates changes in evidence quality, response caution, and bias, enabling mechanistic interpretation of latent decision states. We demonstrate below how the DDM-HMM can identify these parameters, offering a complementary tool for serially-dependent decision data for free-response tasks.

### 5.4 DDM-HMM accounts for speed and accuracy at the individual and population levels

We fit the model to data from 18 animals using an adaptation of the Expectation–Maximization algorithm. To determine how many latent states best captured our data, we computed the difference in model likelihood compared to a classical DDM (a one-state model) on held-out data via five-fold cross validation. Test log-likelihood per trial (ΔLL/trial) averaged across five splits of the data increased monotonically with additional latent states for every animal and plateaued at *K* = 3–4 (**Figure 3A**). Because there was no sharp optimum and gains beyond four states were modest, we selected a four-state model for all subsequent analyses to balance predictive performance and interpretability.^28^ Importantly, improvements relative to simpler models were consistent across animals, indicating that multiple latent states are required to capture structured behavioral variability. Post-hoc analysis of the fit model revealed the DDM-HMM accurately captured individual animal reaction time distributions (**Figure 3B**; **Supplemental Figure 5**) and animal choices. We found a high degree of model agreement between both the quantiles of the aggregated animal RT data and simulated RT data (**Figure 3C**), and the predicted accuracy and the observed accuracy of the animals (**Figure 3D**). These results confirm the DDM-HMM simultaneously accounts for both speed and accuracy at the individual and population levels.

**Figure 3:**
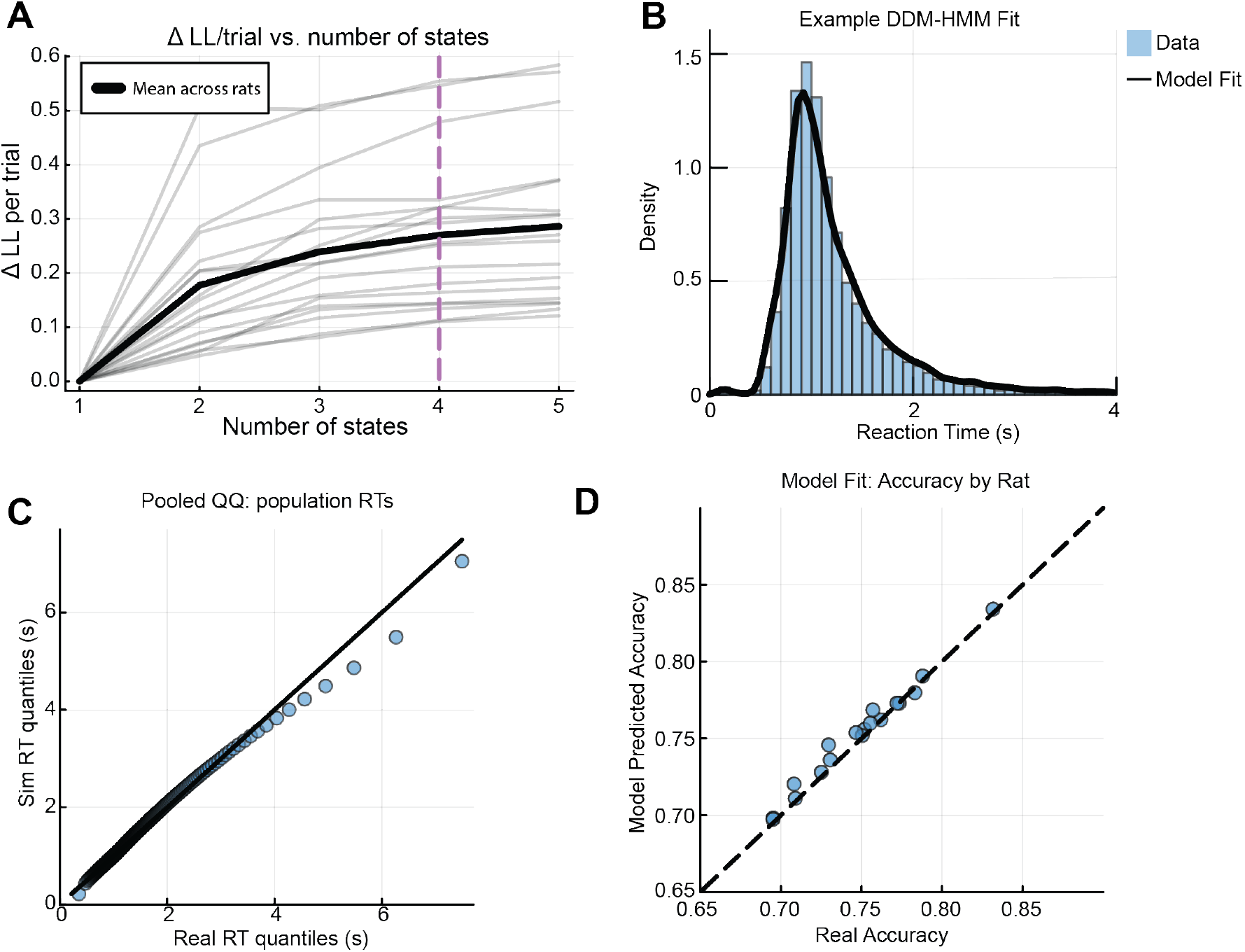
The DDM-HMM provides an accurate account of behavior. **(A)** Results of five-fold cross-validation across animals. The pink dashed line indicates the selected number of states. **(B)** Example model fit of the DDM-HMM (black) to an example rat reaction time (RT) distribution (blue). **(C)** Q–Q plot comparing aggregated RT data to RTs simulated from the DDM-HMM. **(D)** Model-predicted accuracy versus observed animal accuracy.

### 5.5 DDM-HMM reveals discrete, serially-dependent, circadian-modulated decision states

To illustrate the structure of latent decision dynamics identified by the DDM-HMM for single animals, we examined a representative rat (**Figure 4A**). The model identified four latent states with distinct parameter combinations (**Figure 4B**) that systematically related to animal reaction time (**Figure 4C**) and accuracy (**Figure 4D**). One state exhibited high drift rate and an elevated decision boundary and was associated with the highest task performance and longest median reaction time. Although states do not have inherent semantic labels, we referred to this state as a “patient” state given its DDM parameters and resulting behavior. Two states, which we labeled as “impulsive states”, showed intermediate drift rates and lower boundaries, and decreased accuracy and median reaction time. A rare “noise state” (<1% of trials) displayed low drift, high bias, and minimal non-decision time, and lowest overall accuracy and fastest reaction times. The correspondence between DDM-HMM state-wise parameters and accuracy and reaction times demonstrate the utility of the DDM-HMM for partitioning time-varying decision making behavior into unique behavioral strategies. To provide additional support for the model goodness of fit, we examined how simulated reaction times from each model state corresponded with the empirical reaction time distributions (**Figure 4C**). The model provided an accurate generative account of the data; simulated marginal reaction time distributions closely matched the empirical distributions for correct and incorrect trials across states (**Figure 4C**), reproducing the skewed shape and tail structure of the observed data. An important feature of our model is the ability to identify temporal dependence in decision-making behavior across trials. To examine the persistence of DDM states across trials, we computed the posterior state probabilities, which specify the likelihood of the data on each state being generated from each state (**Figure 4E**). Values were typically near 0 or 1, indicating strong confidence of a unique state being expressed on each trial. Furthermore, values exhibited extended dwell times indicating that states persist across many successive trials. This persistence implies that reaction times exhibit serial dependence across trials, a feature of the data that the HMM-DDM can capture and classical decision-making models cannot. Consistent with this prediction, the empirical autocorrelation function (ACF) revealed significant long-timescale structure in RTs (**Figure 4F**). Importantly, the empirical ACF fell within the credibility interval of the model for the first 18 lags, indicating that the DDM–HMM accurately captures these slow temporal dependencies. Beyond serial dependence in decisions, our GAMMs (**Figure 2D–F**) imply that states are more common at specific times of the day. We found DDM-HMM state occupancy was strongly modulated across the 24-hour cycle (**Figure 4G**): the patient state predominated during the lights-off period, whereas impulsive state 1 dominated during the light phase, with impulsive state 2 and the noise state emerging around feeding time. Similar results were obtained after fitting to an expert rat with exceptional performance, indicating that the inferred decision states generalize to highly proficient behavior (**Supplemental Figure 5**).

**Figure 4:**
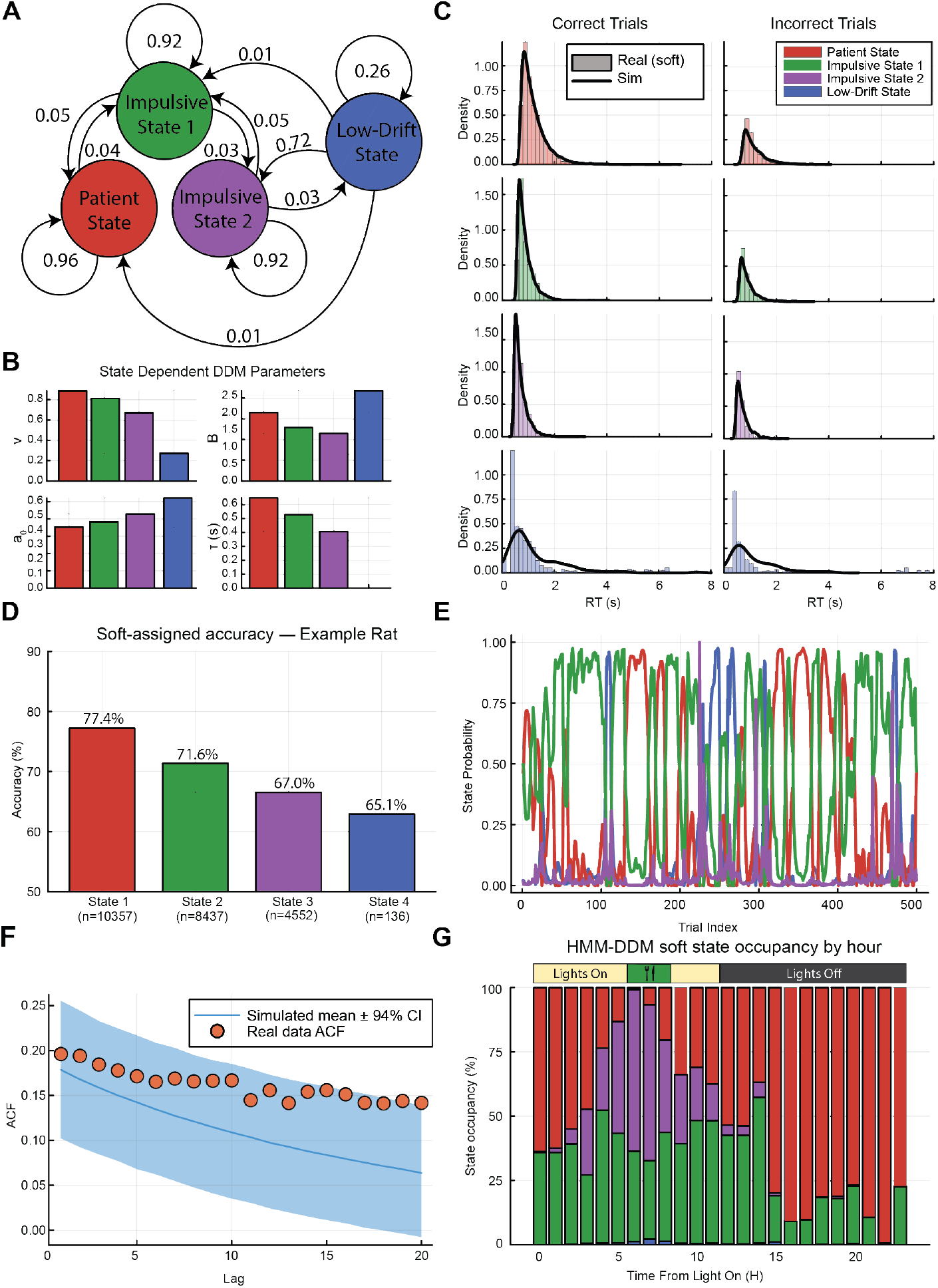
DDM-HMM fit for an individual rat. **(A)** Schematic of the transition dynamics between the four inferred latent decision states: patient, impulsive state 1, impulsive state 2, and low-drift. **(B)** State-dependent drift–diffusion model (DDM) parameters, including drift rate, boundary separation, starting bias, and non-decision time. **(C)** Reaction time (RT) distributions for correct trials shown for each state. Histograms reflect soft-segmented data, and black curves indicate simulated data from the fitted DDM-HMM. **(D)** Soft-assigned choice accuracy for each state. **(E)** Example posterior state probabilities across trials during feeding. **(F)** Autocorrelation function (ACF) of inferred states in the empirical data (points) compared with simulated data from the model (mean *±* 94% CI). **(G)** Hourly state occupancy relative to the light–dark cycle.

### 5.6 Circadian modulation of decision states generalizes across animals

We next evaluated how the DDM-HMM decision making states fluctuated across the 24-hour cycle in the entire population of rats. To compare across animals, we rank-ordered each animal’s latent states by accuracy, from lowest (State 1) to highest (State 4), and computed state occupancy as a function of time of day after rank-ordering states by accuracy within each animal. In hour-long bins, we calculated the posterior probability of occupying each state and averaged these values across animals. We identified a pronounced temporal structure in state occupancy (**Figure 5A**): Low-accuracy states (States 1 and 2) were most prevalent during feeding periods and early light-phase hours, whereas high-accuracy states (States 3 and 4) dominated during the dark cycle.

**Figure 5:**
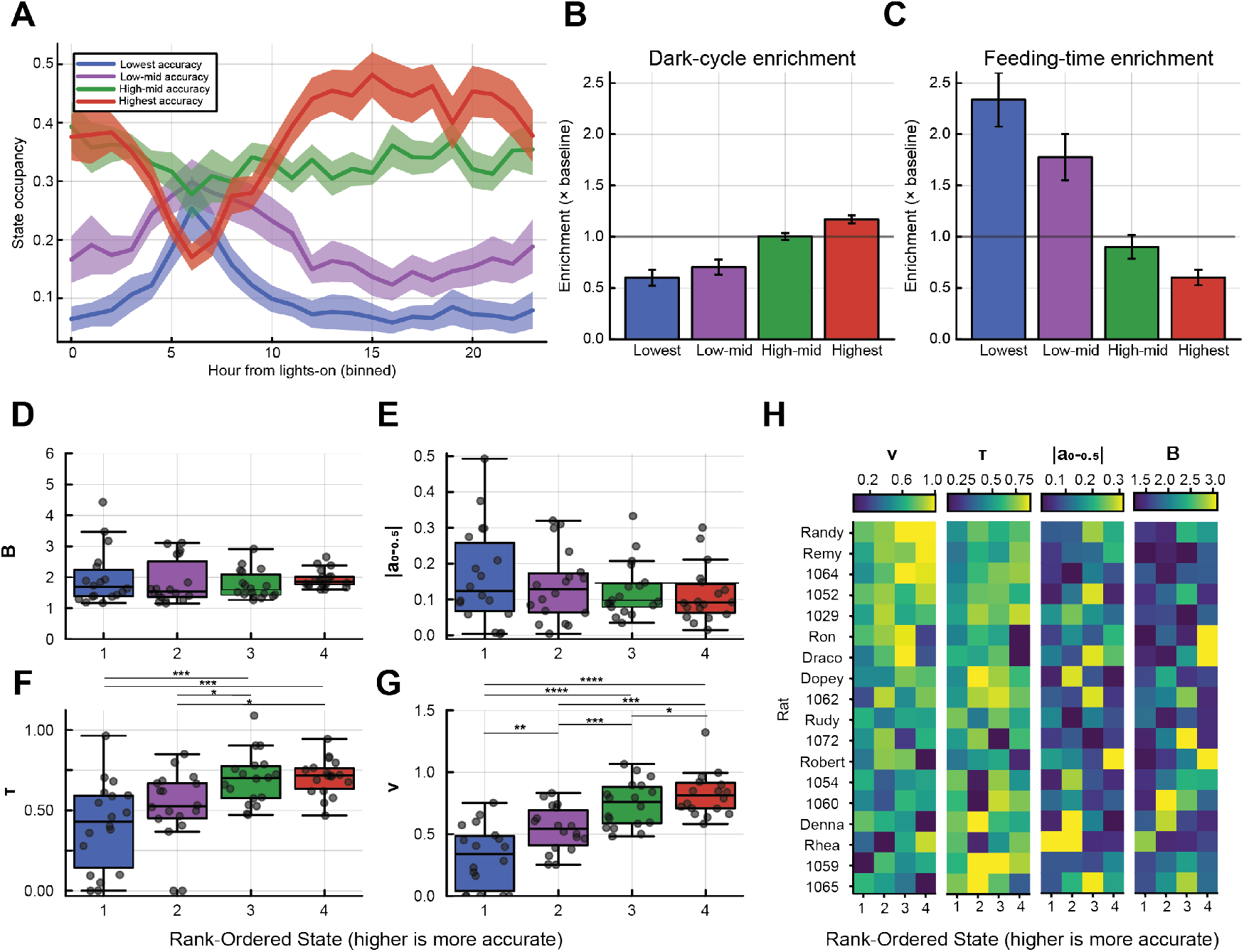
DDM-HMM reveals distinct behavioral states across the 24-hour cycle. **(A)** 24-hour time course of accuracy-ranked state occupancies. **(B)** Enrichment of state occupancy during the dark phase. Lower-accuracy states are less enriched than expected by chance. **(C)** Same as (B), but during feeding. Lower-accuracy states are over-enriched during this period. **(D)** Boundary separation across accuracy-ranked states. **(E)** Starting-point bias across accuracy-ranked states. **(F)** Non-decision time increases with state accuracy. **(G)** Drift rate increases with state accuracy. **(H)** Heat map of the four free DDM parameters. Rats are sorted by mean drift rate (e.g., Denna has the highest average drift rate).

To quantify these effects, we computed enrichment scores for each state during the dark cycle and feeding epochs, defined as the ratio between the probability of occupying a given state during a specific epoch and its overall probability across the session. Low-accuracy states were significantly under-enriched during the dark phase and over-enriched during feeding (**Figure 5A–C**). In contrast, the highest-accuracy state was strongly overrepresented during the dark cycle and suppressed during feeding, while the second-highest accuracy state was expressed at near-chance levels across both epochs. These results indicate that transitions between latent decision states are tightly coupled to the local environment and intrinsic behavioral states.

### 5.7 Distinct combinations of DDM parameters define discrete behavioral states

Having established that accuracy-defined decision states fluctuate across the 24-hour period, we sought to understand how changes in DDM parameters drove these changes in behavior. We found that individual animal state-dependent DDM parameters demonstrated a systematic relationship with state-specific reaction time and accuracy (**Figure 4**). By regressing individual animal state-wise DDM parameters against state-wise accuracy (see Methods), we found drift rate and boundary separation were the dominant predictors of model accuracy (**Supplemental Figure 7A**). In contrast state-wise mean reaction time was largely driven by boundary height and non-decision time (**Supplemental Figure 7C**).

To identify model components that best differentiate states, we examined state-specific DM parameters across animals using a repeated-measures design. For each parameter, we performed a one-way repeated-measures ANOVA with state as the within-subject factor. State-dependent drift rate and non-decision time differed significantly ((*F* (3, 51) = 35.73, *p* = 1.39 *×* 10^−12^, 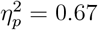 and *F* (3, 51) = 11.14, *p* = 9.80 *×* 10^−6^, 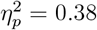 respectively), while we detected no changes in bias magnitude or boundary separation (*F* (3, 51) = 0.77, *p* = 0.52, 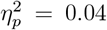 and *F* (3, 51) = 0.35, *p* = 0.78, 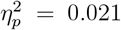, respectively). To characterize specific state differences underlying these effects, we conducted post hoc paired t-tests with Holm correction between all state pairs. Post-hoc comparisons revealed significant drift-rate differences between all tested state pairs: states 3 and 1 (*t* = − 8.43, *p*_*Holm*_ = 9.91 *×* 10^−7^), states 1 and 4 (*t* = 8.36, *p*_*Holm*_ = 9.91 *×* 10^−7^), states 2 and 3 (*t* = 5.01, *p*_*Holm*_ = 4.28 *×* 10^−4^), states 2 and 4 (*t* = 4.55, *p*_*Holm*_ = 8.51 *×* 10^−4^), states 2 and 1 (*t* = − 3.91, *p*_*Holm*_ = 0.00224), and states 3 and 4 (*t* = 2.45, *p*_*Holm*_ = 0.0255; **Fig. 5F–G**), indicating strong and systematic modulation of evidence accumulation across states. Non-decision time differed significantly across four of the six tested state contrasts: states 1 vs. 4 (*t* = 5.02, *p*_*Holm*_ = 6.31 *×* 10^−4^), 3 vs. 1 (*t* = −4.00, *p*_*Holm*_ = 0.00459), 2 vs. 3 (*t* = 3.64, *p*_*Holm*_ = 0.00602), and 2 vs. 4 (*t* = 3.66, *p*_*Holm*_ = 0.00602). Comparisons between states 2 vs. 1 and 3 vs. 4 were not significant after Holm correction (both *p*_*Holm*_ = 0.438; **Fig. 5D–E**).

Although we did not detect significant state-dependent differences in bias magnitude or boundary separation (**Fig. 5G–H**), we asked whether these parameters were necessary to support the full DDM-HMM. To test whether the model was over-parameterized, we performed a BIC-based model comparison in which we refit the DDM-HMM while constraining selected parameters to be shared across states. Specifically, we compared the full DDM-HMM against reduced models with global bias, global non-decision time, global drift rate, global boundary separation, or both global bias and global non-decision time. Each candidate model was fit 30 times from random initializations, and the best-fitting solution was retained for comparison. Across animals, the full DDM-HMM provided the best model fit by BIC, indicating that allowing DDM parameters to vary across latent states was supported despite the absence of significant pairwise effects for some individual parameters (**Supplemental Fig. 8A–B**).

These results demonstrate that fluctuations in decision-making emerge from distinct combinations of DDM parameters—most critically drift rate and non-decision time—that define discrete behavioral states, providing a mechanistic link between parameter-level dynamics and transitions among a set of decision-making regimes.

### 5.8 Ambient light affects drift rate, but not RT, boundary or non decision time

We sought to quantify the effect of changes in ambient illumination on model parameters as it varies with time of day in the live-in system. We trained six rats in the same evidence accumulation task, in a two-hour daily training system. We measured performance on five sessions each with the light on or light off, having the animals perform the task at the same time of day, and fit a one state DDM to each set of sessions. Low ambient light (‘high contrast’) drove a significant change in RT (*t* = 3.010, *p* = 0.030), accuracy (*t* = 6.870, *p <* 0.001), and drift rate (*t* = 6.334, *p* = 0.0014) (**Supplemental Fig. 9A–F**) but had no effect on boundary separation, non-decision time, or starting-point bias (**Supplemental Fig. 9F**). Interestingly, lights-off conditions in the two-hour system were associated with faster reaction times, whereas the high-contrast dark phase in the live-in system was associated with slower reaction times despite higher accuracy. Thus, stimulus contrast (a consequence of different ambient light levels) contributes to the elevated drift rate observed during dark-phase periods in the 24-hour dataset but cannot be reliably linked to changes in other decision parameters or in RT.

### 5.9 The DDM-HMM addresses a central shortcoming of the DDM

To determine whether the structure identified by the DDM-HMM could be explained by established alternatives, we compared its performance to a multilevel drift diffusion model (similar to the “extended DDM”)^29^, which permits trial-to-trial variability in DDM parameters but assumes conditional independence across trials.

The multilevel DDM converged reliably for all animals (**Supplemental Fig. 10A**) and successfully captured marginal distributions of trial-level parameters (**Supplemental Fig. 10B**). To test the model’s appropriateness for the data, we examined if the learned trial-level parameters obey the assumptions of the model. When plotting the auto-correlation function of the learned parameters, we found they exhibited substantial autocorrelation (**Fig. 6A**), violating the model’s assumption that trial-level fluctuations are i.i.d. Furthermore, simulations from the multilevel DDM exhibited near-zero reaction time autocorrelation across lags (**Fig. 6B**), in contrast to the strong serial dependencies in reaction time reflected in the data. These results demonstrate the inadequacy of standard approaches when serial correlation is present in decision-making data, underscoring the shortcoming the DDM-HMM was designed to overcome.

**Figure 6:**
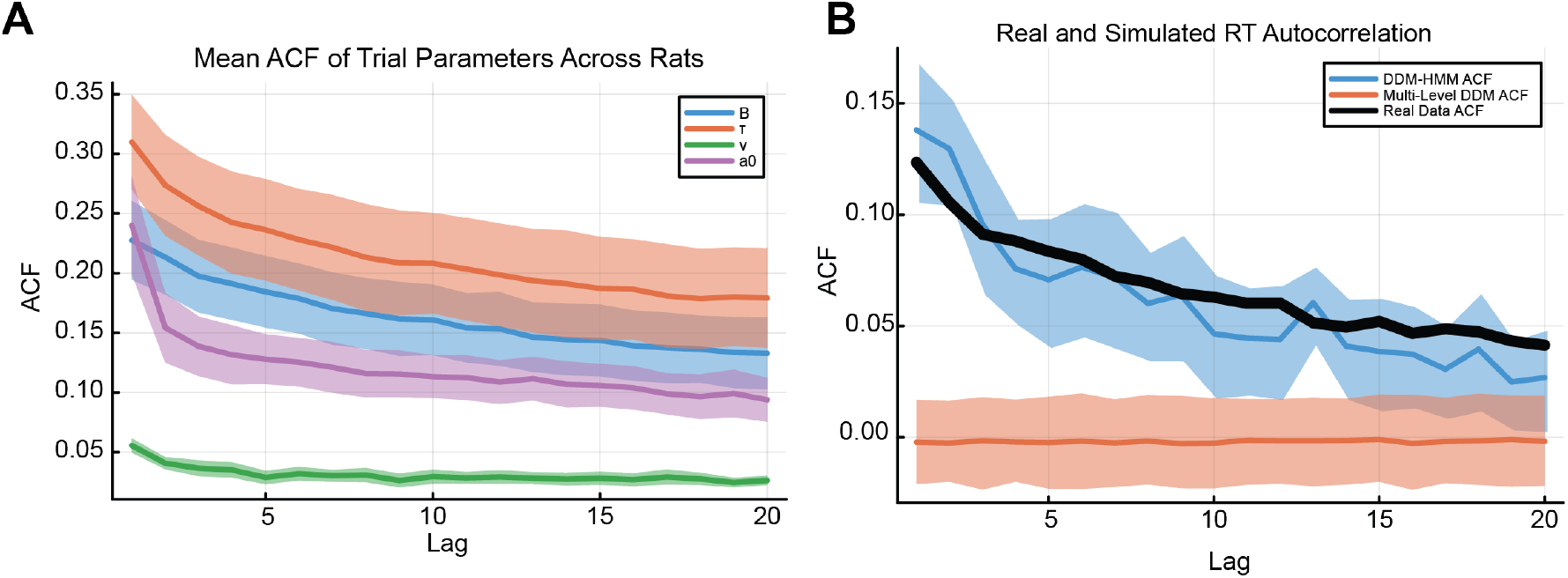
The DDM-HMM captures trial-to-trial correlations in reaction times. **(A)** Mean autocorrelation functions (ACFs) of trial-level DDM parameters estimated by the multilevel DDM, averaged across rats. Shaded regions indicate *±*1 SEM across animals. Under the generative assumptions of the multilevel DDM, trial-level parameters are independent across trials; thus, autocorrelation is expected to be zero at non-zero lags. The observed systematic autocorrelation indicates unmodeled temporal structure. **(B)** Reaction time autocorrelation functions for empirical data (black), simulations from the DDM-HMM (blue), and simulations from the multilevel DDM (orange). Shaded regions indicate 94% credible intervals for simulated data. The DDM-HMM reproduces the serial dependence observed in empirical reaction times, whereas the multilevel DDM fails to capture this structure.

### 5.10 The DDM-HMM discovers similar latent dynamics to the GLM-HMM

A second established alternative for analyzing state-dependent decision making is the GLM-HMM, which models state-dependent choices using a generalized linear model (GLM) and state transitions with an HMM. We fit GLM-HMMs to our data using established approaches.^23,24^ To assess whether both models identified similar latent decision states, we quantified how well aligned the posterior distributions of each model for each animal were using the normalized mutual information (NMI). For all animals, NMI significantly exceeded chance (Benjamini–Hochberg; *FDR* = 0.05; **Supplemental Fig. 11A–B**) indicating the HMM-DDM identifies states similar to those identified by the GLM-HMM. Further, model-based choice predictions were comparable between models indicating that the models had equal potential for modeling choice data (**Supplemental Fig. 11C–D**).

We examined fits in two representative animals (*K* = 3 states). In both cases, the GLM-HMM identified three regimes primarily differentiated by the stimulus gain, consistent with differences in task engagement (**Supplemental Figs. 12A, 13A**). As in the DDM-HMM, these regimes exhibited differences in accuracy and a clear gain–accuracy relationship (**Supplemental Figs. 12B–C, 13B–C**). Both models also produced similar 24-hour state occupancy patterns, supporting the conclusion that animals transition between task-linked behavioral modes (**Figure 4F**; **Supplemental Figs. 11–13E**). The primary qualitative difference concerned state segmentation: although both models were typically confident in state assignment, the GLM-HMM exhibited longer dwell times (**Supplemental Figs. 12D, 13D**), likely reflecting the DDM-HMM’s use of both choice and reaction time to detect transitions, increasing sensitivity to state switching.

### 5.11 The DDM-HMM captures behavioral fluctuations in session-based training

Although the DDM-HMM was motivated by the unique 24-hour continuous dataset we collected, we next asked whether the model generalizes to traditional session based training.^30^ To this end, we fit the model to data from six rats performing the same task during fixed daily two-hour training epochs (**Figure 7**).

**Figure 7:**
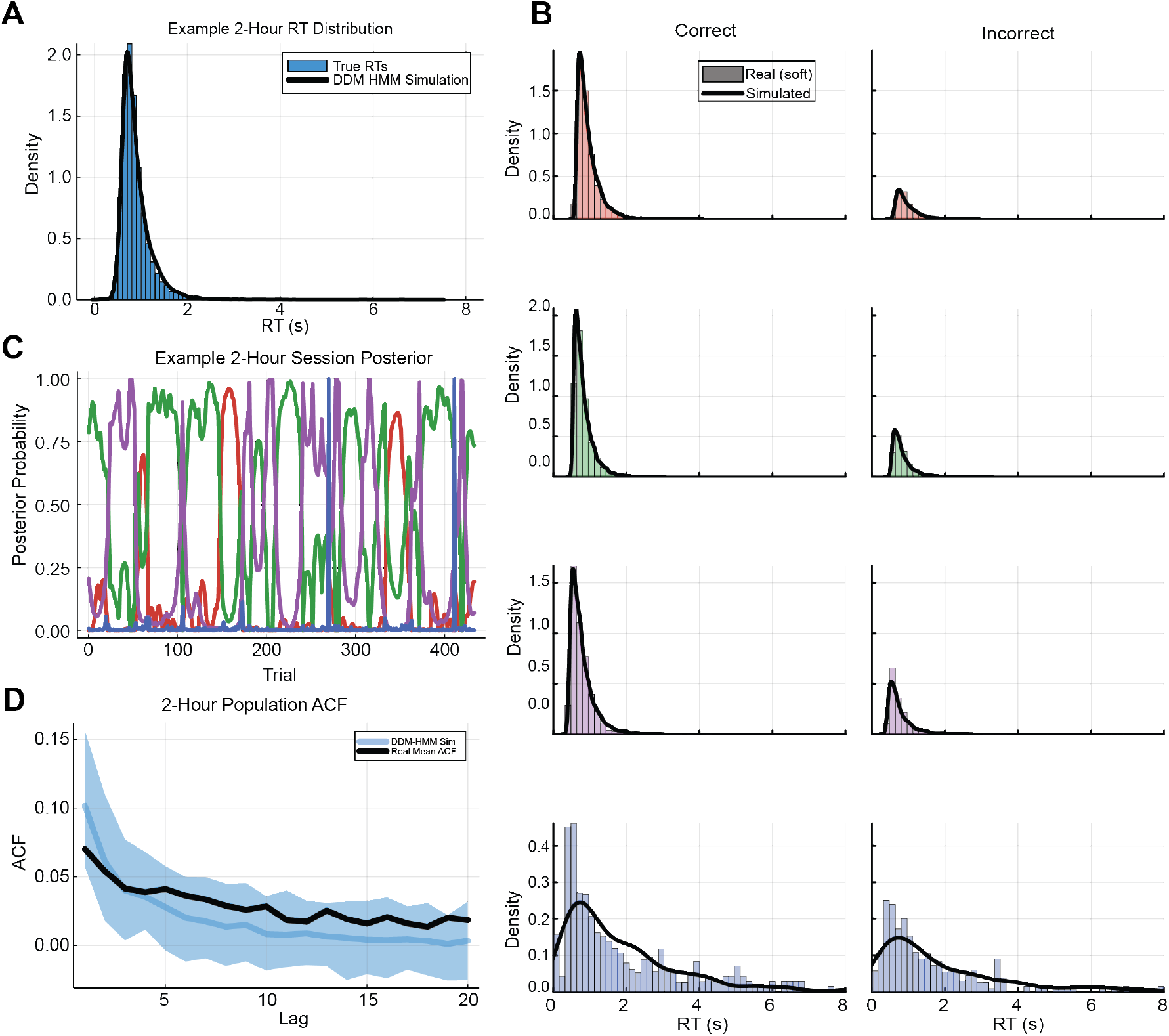
The DDM-HMM captures short-timescale fluctuations in behavior. **(A)** Example reaction time (RT) distribution from a rat performing the task within a fixed 2-hour epoch. **(B)** Learned response time distributions of the corresponding DDM components in the DDM-HMM. **(C)** Example posterior state probabilities over a 2-hour session. **(D)** Population autocorrelation function (ACF) across six rats performing the task in 2-hour epochs. The DDM-HMM population ACF closely matches the empirical ACF.

Within individual two-hour sessions, the DDM-HMM closely reproduced empirical reaction time distributions (**Figure 7A–B**). For a representative rat we found that simulated reaction times from the fitted DDM-HMM matched the full reaction time density (**Figure 7A**), as well as the conditional distributions for correct and incorrect trials (**Figure 7B**), indicating that the inferred state-dependent DDMs captured both accuracy-dependent and error-related RT structure. The model inferred dynamic switching among latent decision states within the session (**Figure 7C**), with posterior state probabilities exhibiting structured transitions across trials. Importantly, when aggregating across animals, simulations from the DDM-HMM reproduced the empirical reaction time autocorrelation function within two-hour blocks (**Figure 7D**), capturing the magnitude and decay profile of serial dependence.

These results demonstrate that the DDM-HMM does not merely explain slow, circadian-scale modulation, but also captures structured trial-to-trial dynamics on shorter timescales. Together with the cross-model comparisons, this establishes that incorporating both reaction times and explicit state transitions is necessary to explain the temporal organization of decision-making behavior across timescales.

## 6 DISCUSSION

We examined how perceptual decision-making varies across the 24-hour cycle by combining continuous, autonomous behavioral measurement with a generative modeling framework designed to capture long-timescale fluctuations in behavior. By allowing rats to perform trials continuously over 24-hour periods, for weeks to months, we observed pronounced and reliable fluctuations in accuracy, reaction time, and task engagement aligned with both the light–dark cycle and feeding schedule. These effects were robust at the population level yet expressed with substantial inter-animal variability, underscoring a central point: decision-making policies are not static traits measured in isolated sessions, but dynamically regulated processes that evolve across biologically meaningful timescales.

A central contribution of this work is to formalize this non-stationarity as structured switching between discrete decision-making strategies. Existing modeling approaches capture only part of this structure. Classical drift diffusion models provide mechanistic interpretability but assume that all trials arise from a single, time-invariant process. Multilevel extensions relax parameter constancy but retain conditional independence across trials, rendering them incapable of generating the sustained reaction-time autocorrelations observed here. State-switching models such as the GLM-HMM can identify discrete behavioral modes but operate on choices alone, leaving the cognitive parameters underlying reaction-time distributions unmodeled.

The DDM-HMM integrates these two traditions. By embedding state transitions within a mechanistic evidence-accumulation model, it captures both the full reaction-time distribution and the persistent serial dependencies that characterize long-duration behavior. Critically, the model does not merely fit marginal statistics: it reproduces the empirical autocorrelation structure of reaction times across multiple lags, a feature that alternative DDM-based approaches fail to explain. This capacity to jointly account for distributional form and temporal dependence is essential, because reaction times disambiguate changes in sensory gain, response caution, and bias—distinctions that are behaviorally indistinguishable at the level of choices alone. Incorporating reaction-time dynamics is not a modeling luxury but a necessity for mechanistic inference in non-stationary decision-making.

The mechanisms underlying these state transitions remain to be determined. One possibility is sensory adaptation: as nocturnal animals with rod-dominated retinas, rats may experience reduced effective contrast under increased ambient illumination, leading to diminished sensory gain.^31–34^ Alternatively, light-dependent modulation of arousal and neuromodulatory tone could alter both evidence accumulation and non-decisional processes.^10,12,17,35^ A recent mouse study reported higher cFos activity in reward-related regions during the subjective night.^36^ Increased activity in reward circuits may reflect enhanced motivation, consistent with the higher trial rate we observed during the subjective night. Disentangling these mechanisms will require targeted manipulations of sensory, arousal, and motivational variables.

Finally, motivational dynamics associated with scheduled feeding likely contribute to state transitions. Feeding anticipatory activity (FAA) has been widely documented in food-restricted animals and can occur independently of canonical circadian oscillators.^37^ In our data, feeding epochs were associated with increased trial initiation in some animals but generally reduced accuracy and shortened reaction times, suggesting a shift toward less deliberative responding. These effects are compatible with motivationally driven adjustments in urgency or engagement. These findings extend prior work linking circadian and feeding dynamics to behavioral performance by demonstrating that such factors reshape the computational structure of perceptual decision-making. Rather than reflecting stable traits, decision policies fluctuate across biologically relevant timescales. Accordingly, experimental designs and modeling approaches that assume stationarity risk conflating intrinsic computational differences with contextual modulation. Careful consideration of animals’ circadian biology and motivational state is therefore essential for accurate interpretation of decision-making behavior.

### 6.1 Limitations of the study

Despite its strengths, several limitations warrant consideration. First, although the DDM-HMM identifies persistent latent decision states, it does not specify the neural or physiological mechanisms governing transitions between them. Linking these computational regimes to biological substrates will require future studies combining continuous behavioral monitoring with neural recordings or targeted perturbations. The standard probabilistic structure of the model can easily accommodate available neural measurements to better identify latent cognitive states, making it a powerful general purpose tool for future studies.

Second, we interpreted drift-rate changes as reflecting variation in effective evidence quality, but this parameter aggregates multiple processes, including sensory encoding, attention, and internal noise. Our controlled illumination experiment demonstrated that ambient light selectively modulates drift rate, implicating stimulus contrast as one contributor, though the opposing reaction time effects between the two-hour and live-in systems (see Results) suggest that additional factors beyond illumination — such as arousal or circadian phase — also shape decision dynamics. Disentangling these components will require task designs that independently manipulate stimulus strength and motivational or attentional demands, as well as modeling approaches that explicitly separate sensory and cognitive contributions.

Third, aspects of the task design introduce potential confounds. Feeding required manual intervention, increasing experimenter presence during the light phase. Although animals were highly trained and habituated, we cannot exclude the possibility that human interaction contributed to some observed effects. Future implementations incorporating automated feeding could further reduce this source of variability.

Finally, limitations inherent to the drift diffusion framework remain. Parameters such as non-decision time are weakly identifiable and constrained primarily by the leading edge of the reaction-time distribution, and interpretation of changes in this parameter is inherently ambiguous.^38^ Trade-offs between parameters (e.g., drift rate and boundary separation) can also complicate inference, although the large datasets collected here mitigate these concerns.^38^

## 7 METHODS

### 7.1 Experimental Methods

#### 7.1.1 Animal Subjects

All experiments and procedures were performed in accordance with protocols approved by the Boston University Institutional Animal Care and Use Committee (IACUC). A total of 18 Long–Evans rats (12 male, 6 female), aged 3–12 months, were used in this study. Rats were food restricted and maintained above 80% of their baseline body weight throughout training and testing. Animals had *ad libitum* access to water and were housed on a 12 h light/dark cycle, with lights on from 7:30 AM to 7:30 PM.

#### 7.1.2 Automated Operant Training System

Behavioral tasks were administered in custom-built acrylic operant chambers equipped with three nose ports. Each port (Sanworks or custom-built) contained a white LED for visual stimulus presentation, an infrared (IR) LED and photodetector pair for nose-poke detection, and a peristaltic pump for liquid reward delivery. Behavioral control software was written in MATLAB and executed on a Teensy-based microcontroller system (Bpod; Sanworks), which implemented trial-by-trial task logic using a finite state machine architecture.

To enable high-throughput behavioral experiments, we developed BpodAcademy, a Python-based graphical user interface for centralized and simultaneous control of multiple Bpod rigs (https://github.com/RatAcad/BpodAcademy), built on a modified Bpod library (https://github.com/RatAcad/Bpod_Gen2). BpodAcademy communicates with MATLAB to execute state machine–based task protocols that define trial structure, training contingencies, and stimulus–respons logic for each behavioral session. Animals interacted with the system via nose-poke inputs, while task events such as LED cues and liquid reward delivery were controlled as outputs of the state machine. At the conclusion of each session, Bpod generated a .mat file containing trial-by-trial behavioral events and metadata, organized by animal and protocol name and saved locally on the acquisition computer. Behavioral data were automatically transferred nightly to the Boston University Engineering network drive for centralized storage and backup. Data were subsequently ingested into a DataJoint-based pipeline for organization and preprocessing. Raw `.mat` files were parsed to compute summary metrics and task-relevant features, which were flattened into relational tables defined by a shared schema (e.g., animal, protocol, session). The schema is maintained on the lab’s GitHub repository (https://github.com/RatAcad/dj_ratacad) and can be visualized using the DataJoint library. All processed tables were stored in a MySQL database hosted on an AWS server managed by Boston University IT. Researchers accessed the dataset directly from the AWS server for downstream analyses using the lab’s Python interface (`dj_ratacad`) or standard SQL queries.

#### 7.1.3 Live-In training Facility

To enable continuous assessment of behavior across the circadian cycle, we developed RatAcad (Rat Academy), a live-in operant training facility in which rats were housed directly within integrated operant chambers and could engage with the task freely over a full 24-hour period (**Figure 1A**). Custom plexiglass enclosures incorporated the behavioral apparatus while permitting visual and olfactory contact with conspecifics. Animals had continuous access to water and received daily pellet delivery, allowing task engagement at any time of day without water restriction, which is healthier for the animals.^39^

Task parameters remained constant throughout experiments: the light–dark cycle (7:30 am lights on; 7:30 pm lights off, EST) and feeding schedule (2:00–4:00 pm daily) were fixed, and animals could initiate trials ad libitum across the circadian cycle. Continuous behavioral monitoring was maintained via Bpod, with automated daily synchronization of data to the central database. Human interaction was limited to essential husbandry procedures, including daily feeding, weekly weighing, and biweekly cage changes performed by Boston University animal care staff, minimizing potential experimenter-induced influences on behavior.

#### 7.1.4 Visual Evidence Accumulation Task

Rats were housed and tested in a customized operant chamber. Each chamber contained three nose ports, each equipped with a white LED. The task has been described previously.^26^ In the final stage of training, visual flashes were presented according to a Bernoulli process such that, on each time step, a flash occurred on the correct side with 75% probability and on the incorrect side with 25% probability. The correct side (left or right) was selected randomly on each trial (**Supplemental Figure 1**). Rats could report their choice at any time during the trial by poking into either side port. Correct responses were rewarded with 25 µl of 10% sucrose, followed by a 5 s inter-trial interval. Incorrect responses triggered a 3 s timeout, resulting in a total 8 s inter-trial interval. If a rat failed to respond within 8 s of trial initiation, the trial was scored as an omission.

#### 7.1.5 Training Pipeline

Rats were single-housed in the operant chambers at the Boston University Animal Science Center and progressed through staged training of a visual evidence accumulation task. Training duration ranged from 1–4 weeks depending on individual performance. Reward volume remained constant throughout training (0.025 mL of 10% sucrose).

Animals completed three core training stages before reaching the final task configuration. In Stage 1 (1–3 days), rats were rewarded for nose-poking into an illuminated side port. In Stage 2 (1–21 days), rats were required to nose-poke into an illuminated center port followed by an illuminated side port to receive reward. In Stage 3, rats nose-poked into the illuminated center port followed by a flashing side port. Once animals achieved >90% accuracy in selecting the flashing port, the relative flash probabilities were progressively adjusted across stages (100:0 → 90:10 → 80:20). After meeting criterion at each condition (typically 1–2 days per condition), rats advanced to the final task configuration with a 75:25 flash probability ratio.

#### 7.1.6 Circadian Housing and Husbandry Conditions

Upon reaching final task form, rats remained housed within the live-in operant chambers for continuous behavioral monitoring. Animals were food-restricted and weighed weekly to maintain body weight above 80% of baseline. During the daily feeding window (2:00–4:00 pm), each rat received three pellets (15 g total chow). Water was available ad libitum. Cage changes were performed biweekly by Boston University animal husbandry staff.

#### 7.1.7 Session based daily training

12 Long–Evans rats were used for the daily training experiments. These animals were housed in a separate room, apart from the live-in training facility. They were transported to the training room for scheduled sessions lasting 2 hours per day, Monday through Friday. Animals were food restricted and were given 15 grams of chow daily after training. Six of the twelve animals participated in the high-contrast and low-contrast control experiments. Each animal completed five high-contrast sessions and five low-contrast sessions, with conditions interleaved across days. During high-contrast sessions, the room was completely dark. During low-contrast sessions, the room lights remained on.

### 7.2 Quantification and statistical analysis

#### 7.2.1 Generalized Additive Models

Generalized additive models (GAMs) extend generalized linear models by replacing fixed linear coefficients with flexible smooth functions of the predictors, letting the data shape each effect rather than imposing a specific parametric form (e.g., sinusoidal). Each smooth is built from a weighted sum of basis functions—here, cyclic cubic regression splines, which enforce wrap-around continuity over the 24-hour cycle—and the wiggliness of the resulting curve is regularized by a smoothing penalty selected from the data via restricted maximum likelihood (REML). Generalized additive *mixed* models (GAMMs) further include random effects, which we used to estimate animal-specific intercepts and animal-specific deviations from the population smooth. Conceptually, this allows us to recover a population-average circadian profile while preserving individual variability, without committing in advance to the shape of the underlying time-of-day modulation.

To characterize within-day changes in behavior while allowing for between-animal variability, we fit generalized additive mixed models (GAMMs) using the `mgcv` package (v1.9-4)^40^ in R^41^ (v4.5.2). Time of day was coded as hours since lights-on,

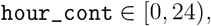

computed from the trial timestamp and wrapped modulo 24 to respect the circadian cycle.

Population-level time-of-day effects were modeled using cyclic cubic regression splines over `hour_cont`. Between-animal variability was captured via random intercepts and factor–smooth interactions, allowing each animal to deviate smoothly from the population-level circadian profile. Smooths were estimated using fast restricted maximum likelihood (REML).

We used moderate basis dimensions for population-level smooths (typically *k* = 16–20) and smaller basis dimensions for animal-specific deviation smooths (typically *k* = 8), together with a mild over-penalization parameter (*γ* = 1.3) to encourage unnecessary smooth components to shrink toward zero and reduce overfitting. All time-of-day smooths were constrained to be cyclic by specifying knots at 0 and 24 hours, ensuring continuity and smoothness at the lights-on/lights-off boundary.

Population-average (“marginal”) curves were obtained by predicting the fitted model for each animal across a dense grid of time-of-day values and averaging predictions across animals at each time point. Uncertainty bands were computed using a nonparametric bootstrap over animals. For each of 300 bootstrap replicates, animals were resampled with replacement, population-average curves were recomputed, and the 2.5th and 97.5th percentiles of the bootstrap distribution at each time point were taken as an approximate 95% confidence interval.

#### 7.2.2 Accuracy Model

Accuracy was modeled at the single-trial level as a Bernoulli outcome indicating whether the response was correct,

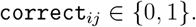

For trial *j* from animal *i* at time of day *t*_*ij*_ (hours since lights-on), we assumed

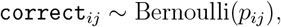

with a logit link and additive structure

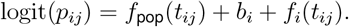

Here, *f*_pop_(*t*) is a cyclic smooth capturing the population-level modulation of accuracy across the day, *b*_*i*_ is an animal-specific random intercept, and *f*_*i*_(*t*) is an animal-specific smooth deviation from the population curve implemented as a factor–smooth interaction. This structure estimates a shared circadian accuracy profile while allowing animals to differ in both overall performance and the shape of their daily modulation. Population-average accuracy across the 24-hour cycle derived from this model is shown in **Figure 2D**; per-animal fits are shown in **Supplemental Figure 3**.

#### 7.2.3 Reaction Time Model

Reaction times were modeled on a continuous scale in seconds using a Gamma distribution with a log link, enforcing positivity and accommodating right-skewed distributions. For trial *j* from animal *i* at time *t*_*ij*_, we assumed

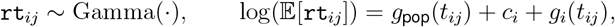

where *g*_pop_(*t*) is a cyclic population-level smooth of time of day, *c*_*i*_ is an animal-specific random intercept, and *g*_*i*_(*t*) is an animal-specific deviation smooth. This model yields a population-level circadian reaction-time profile while allowing individual differences in overall speed and time-of-day modulation. Population-average reaction times across the 24-hour cycle derived from this model are shown in **Figure 2E**; per-animal fits are shown in **Supplemental Figure 2**.

#### 7.2.4 Trial Production (Trial Rate) Model

To model how trial production varied across the 24-hour cycle, behavior was aggregated into regular time-of-day bins within each session. Trials were assigned to bins of width Δ*t* = 10 minutes (Δ*t* = 1*/*6 hours). For each animal *i*, time-of-day bin *b*, and session *s*, we counted the number of trials *Y*_*ibs*_ and defined the exposure as

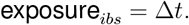

We then collapsed across sessions to obtain, for each animal *i* and bin *b*,

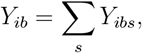

the number of contributing sessions *n*_sess,*ib*_, and the total exposure

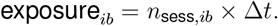

Bins with zero exposure were excluded.

Aggregated counts were modeled using a negative binomial GAMM with a log link and an offset for exposure:

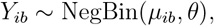

with

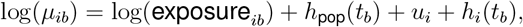

where *t*_*b*_ is the center of bin *b* (hours since lights-on, wrapped to [0, 24)), *h*_pop_(*t*) is a cyclic population-level smooth capturing circadian modulation of trial rate, *u*_*i*_ is an animal-specific random intercept, and *h*_*i*_(*t*) is an animal-specific deviation smooth. The offset ensures that

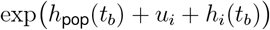

can be interpreted as the expected trial rate (trials per hour) for animal *i* at time of day *t*_*b*_. Population-average trial rates across the 24-hour cycle derived from this model are shown in **Figure 2F**; per-animal fits are shown in **Supplemental Figure 4**.

### 7.3 Drift Diffusion Models

#### 7.3.1 Drift Diffusion Model Implementation

We implemented a standard two-boundary drift diffusion model (DDM) to generate and evaluate trial-wise reaction times and choices. The latent decision variable *a*(*t*) evolves according to the stochastic differential equation

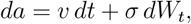

where *v* is the drift rate, *σ* is the diffusion scale, and *W*_*t*_ is a Wiener process. We fixed *σ* = 1, so the noise increment is 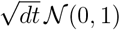.

Decisions occur when the process reaches one of two absorbing boundaries at 0 (lower) and *B* (upper), with *B >* 0 denoting the boundary separation. The starting point is

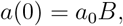

where *a*_0_ ∈ [0, 1] parameterizes the initial bias as a fraction of the boundary separation. Observed reaction time includes a non-decision component *τ*, such that the decision process begins after *τ* seconds.

#### 7.3.2 Stimulus Coding and Choice Convention

Each trial includes a binary stimulus label *s* ∈ {−1, +1}, where −1 denotes leftward evidence and +1 denotes rightward evidence. The drift parameter *v* is treated as a magnitude, and the signed drift on each trial is

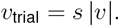

Choices are coded as `choice` ∈ {−1, +1}, where −1 indicates hitting the lower boundary (left) and +1 indicates hitting the upper boundary (right).

#### 7.3.3 Parameter Estimation

Model parameters were estimated via maximum likelihood. Specifically, we solved

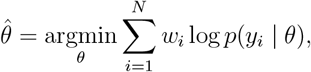

where *y*_*i*_ = (`rt`_*i*_, `choice`_*i*_), *θ* = (*v, B, a*_0_, *τ*), and *w*_*i*_ are optional trial weights (used, for example, in HMM extensions). Trial-wise likelihoods were evaluated using a standard numerical solution to the DDM first-passage-time density.^42^

#### 7.3.4 DDM fitting for the contrast-manipulation dataset

For the contrast-manipulation dataset, DDMs were fit using the `rddm` R package (https://github.com/gkane26/rddm), which estimates DDM parameters via Quantile Maximum Probability Estimation (QMPE).^43^ Under QMPE, the empirical reaction-time distribution for each response is summarized by a small set of quantiles, and parameters are estimated by maximizing a multinomial likelihood over the proportion of trials falling between successive quantile bins as predicted by the DDM first-passage-time density. This quantile-based likelihood is robust to contaminant reaction times and to deviations from the model in the extreme tails of the RT distribution, which is well-suited to the relatively small per-session trial counts in the daily two-hour training data. We fit the standard two-boundary DDM with free parameters (*v, B, a*_0_, *τ*) separately for low- and high-contrast sessions for each animal, and used the resulting fits to generate the psychometric functions, RT distributions, and parameter comparisons shown in **Supplemental Figure 9**.

#### 7.3.5 Multilevel Drift Diffusion Model

As an alternative to state-space models, we fit a multilevel drift diffusion model that allows trial-to-trial variability in DDM parameters while assuming conditional independence across trials given trial-specific parameters. This model treats each trial as having its own latent DDM parameters drawn from a shared Gaussian hyperdistribution, and performs approximate Bayesian inference using variational inference (VI).

#### 7.3.6 Trial-level latent parameters and constrained transforms

For each trial *t*, we introduce an unconstrained latent parameter vector

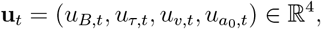

which is mapped to constrained DDM parameters via the deterministic transform used in our implementation:

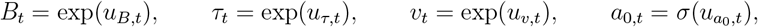

where *σ*(·) denotes the logistic sigmoid and enforces *a*_0,*t*_ (0, ∈ 1). Thus each trial is generated by a standard two-boundary DDM emission likelihood

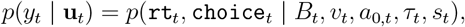

evaluated using the first-passage-time likelihood described above.^42^

#### 7.3.7 Hierarchical prior over trial parameters

Trial-level unconstrained parameters are drawn independently from a diagonal-covariance Gaussian hyperdistribution,

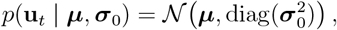

where ***µ*** ∈ ℝ^4^ and 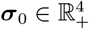 are dataset-level hyperparameters.

#### 7.3.8 Variational approximation and ELBO

We approximated the posterior over each **u**_*t*_ with a diagonal-covariance Gaussian variational distribution,

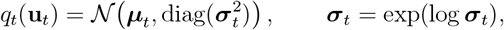

with trial-specific variational parameters {***µ***_*t*_, log ***σ***_*t*_}.

We optimized the evidence lower bound (ELBO)

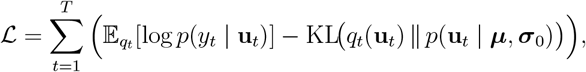

where the KL divergence between two diagonal Gaussians is available in closed form:

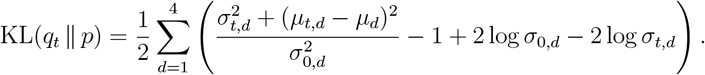

The expected log-likelihood term was estimated using Monte Carlo samples obtained via the reparameterization trick:

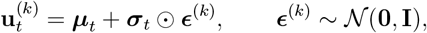

so that

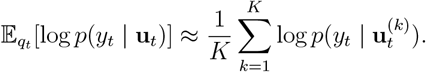

### 7.4 Optimization and coordinate-ascent updates

We optimized the VI objective using a coordinate-ascent scheme alternating between (i) trial-level variational updates and (ii) hyperparameter updates.

#### Trial-level updates

For fixed hyperparameters (***µ, σ***_0_), each *q*_*t*_ was updated by minimizing the negative ELBO for that trial with respect to (***µ***_*t*_, log ***σ***_*t*_). Optimization used BFGS with back-tracking line search (Optim.jl), and gradients were obtained via forward-mode automatic differentiation. To reduce Monte Carlo noise during optimization, the same set of *K* standard-normal samples 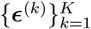 was used within each trial update.

#### Hyperparameter updates

Given updated 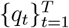, hyperparameters were updated by moment matching. Let *µ*_*t,d*_ and 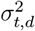 denote the variational mean and variance for trial *t* and dimension *d*.

The hyper-mean was updated as

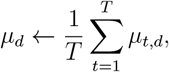

and the hyper-variance was updated as the average of the second moment under *q*_*t*_,

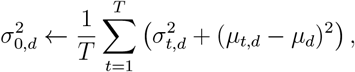

followed by 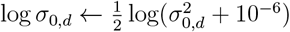 for numerical stability.

#### 7.4.1 Initialization and convergence monitoring

Hyperparameters were initialized using simple data-driven heuristics. Specifically, *τ* was initialized using a low quantile of reaction times (10th percentile), *B* using the reaction-time standard deviation scaled by a constant factor, and *v* using a logit-like transform of empirical accuracy, with bounds applied to ensure reasonable initial values. The starting point was initialized as unbiased (*a*_0_ = 0.5). Initial hyperparameter standard deviations were set to relatively broad values to allow flexibility during inference.

Trial-level variational parameters were initialized by perturbing the hyper-mean with small Gaussian noise and setting trial-level log-standard deviations equal to the initial hyper log-standard deviations. We tracked the total ELBO across VI iterations as a convergence diagnostic. ELBO trajectories and inferred trial-level hyperparameter posteriors for the multilevel DDM are shown in **Supplemental Figure 10**; comparisons of its predicted reaction-time autocorrelations to those of the DDM–HMM and the empirical data are shown in **Figure 6**.

### 7.5 Hidden Markov Models

We modeled trial-by-trial behavior using hidden Markov models (HMMs), in which observed data are generated by an unobserved discrete latent state that evolves according to a first-order Markov process. Let *z*_*t*_ ∈ {1, …, *K*} denote the latent state on trial *t*. The latent state dynamics are defined by an initial distribution ***π*** and a transition matrix **A**:

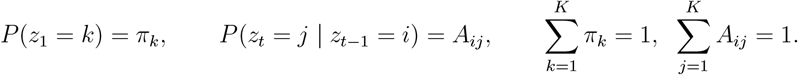

Conditional on *z*_*t*_, observations *y*_*t*_ are generated independently according to a state-specific emission model,

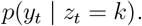

#### 7.5.1 MAP estimation with Dirichlet priors

To regularize state occupancy and transitions, we placed Dirichlet priors on ***π*** and on each row of **A**:

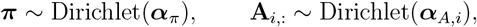

where 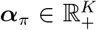 and 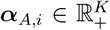 (one vector per row) are strictly positive concentration parameters. These priors yield maximum-a-posteriori (MAP) updates in the Baum–Welch algorithm by contributing pseudo-counts (*α* − 1) in the M-step.

Parameters were estimated via EM. In the E-step, posterior state marginals

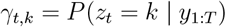

and expected transition counts

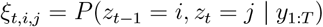

were computed using the forward–backward algorithm. In the M-step, the MAP updates for ***π*** and **A** were

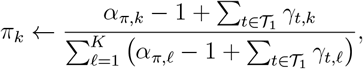

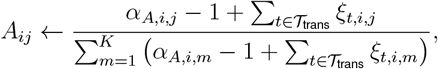

where 𝒯_1_ indexes the first trial of each session (or sequence) and 𝒯_trans_ indexes transitions within sequences. Emission parameters were updated using the posterior state probabilities as trial weights (responsibilities), i.e.,

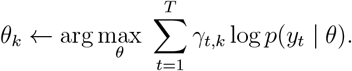

#### 7.5.2 Direct likelihood optimization via L-BFGS

For models where the M-step of Baum–Welch is analytically tractable, we used the MAP-EM procedure described above. However, some emission distributions—including the first-passage-time density of the DDM—do not admit closed-form M-step updates, and we found in practice that Baum–Welch occasionally failed to converge to a competitive local optimum for the DDM– HMM, particularly under parameter constraints (see below). In these cases, we bypassed EM and directly maximized the marginal log-likelihood of the full model,

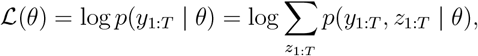

which can be evaluated in *O*(*K*^2^*T*) time via the forward algorithm. Gradients with respect to the full parameter vector (initial distribution, transition matrix, and emission parameters) were obtained by forward-mode automatic differentiation through the forward recursion using ForwardDiff.jl^44^, and optimization was performed using the L-BFGS algorithm in Optim.jl^45^. Stochastic parameters were optimized through a softmax reparameterization, and strictly positive parameters were optimized in log-space to remove inequality constraints. This procedure was used as an alternative or fallback when EM did not converge reliably.

### 7.6 Drift Diffusion Model–Hidden Markov Model (DDM–HMM)

In the DDM–HMM, each latent state *k* is associated with a distinct drift diffusion model (DDM) that generates both reaction times and choices. Conditional on *z*_*t*_ = *k*, trial *t* is generated by a DDM with state-specific parameters

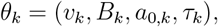

and emission likelihood

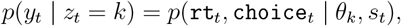

evaluated using the first-passage-time density of the Wiener process.^42^

We estimated parameters using MAP Baum–Welch with Dirichlet priors on the initial distribution and transition matrix, as described above. In the M-step, state-specific DDM parameters were updated by maximizing the responsibility-weighted DDM log-likelihood:

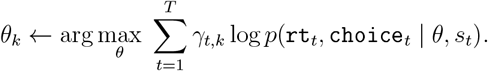

To optimize this function, we used the L-BFGS algorithm included in Optim.jl^45^ (v1.13.2). Gradients were computed using forward-mode automatic differentiation via the ForwardDiff.jl package^44^ (v1.3.0). Cross-validated selection of the number of states and overall model-fit diagnostics for the DDM–HMM are shown in **Figure 3**; state-specific DDM parameters and per-state behavioral summaries for an individual rat are shown in **Figure 4**; population-level state occupancy across the 24-hour cycle, together with statistical comparisons of state-specific parameters, are shown in **Figure 5**.

#### 7.6.1 Constrained-parameter model comparison via BIC

To test whether each state-specific DDM parameter was necessary to explain the data, we compared the full DDM–HMM against a set of reduced models in which selected DDM parameters were constrained to be shared across latent states. We fit five reduced models, in each of which one of the following was tied to a single global value across states: drift rate, boundary separation, starting-point bias, non-decision time, or jointly bias and non-decision time. All non-constrained parameters were still allowed to vary across states.

To accommodate the rugged DDM–HMM likelihood landscape and reduce sensitivity to initialization, each candidate model was fit 30 times from random initializations. Initial values were drawn by adding Gaussian perturbations to the parameter estimates of a single-state DDM fit on the same data, providing a behaviorally plausible baseline from which random initializations could explore the landscape. Across the 30 fits, the solution with the highest marginal loglikelihood was retained. For models in which the constrained M-step was awkward, fits were obtained using the direct-likelihood L-BFGS procedure described above.

Models were compared by computing the Bayesian Information Criterion (BIC),

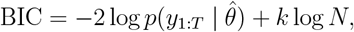

where *k* is the number of free parameters in the model (initial distribution, transition matrix, and unconstrained emission parameters) and *N* is the total number of trials. Lower BIC indicates a better fit after penalizing model complexity. Results of this analysis are shown in **Supplemental Fig. 8A–B**.

### 7.7 GLM–HMM Models

In the GLM–HMM, the trial-wise choices were modeled using a Bernoulli logistic regression observation model within each latent state. Let *y*_*t*_ = 1 denote a rightward choice on trial *t*. Conditional on latent state *z*_*t*_ = *k*, the choice probability was

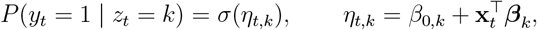

where *σ*(·) is the logistic sigmoid function, *β*_0,*k*_ is a state-specific intercept, and ***β***_*k*_ are statespecific regression coefficients.

The covariate vector **x**_*t*_ included (i) flash ratio, defined as (*N*_*R*_ − *N*_*L*_)*/*(*N*_*R*_ + *N*_*L*_), capturing sensory evidence strength and direction; (ii) previous choice; and (iii) previous reward. Positive coefficients indicate increased probability of choosing the right port.

To reduce overfitting and improve numerical stability, all non-intercept regression coefficients were regularized with an 𝓁_2_ penalty, equivalent to a zero-mean Gaussian prior,

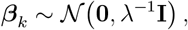

where *λ* controls the strength of regularization. Parameters were estimated using responsibility-weighted logistic regression, with posterior state probabilities providing trial-wise weights. Inferred GLM-HMM state structure, psychometric weights, and circadian state occupancy are shown in **Supplemental Figure 12**; an independent replication in an expert rat is shown in **Supplemental Figure 13**.

### 7.8 Bayesian Regression Models

We modeled trial-wise accuracy proportions (or bounded accuracy scores) acc_*i*_ ∈ (0, 1) as a function of standardized DDM-derived predictors. Predictors were formed from (*v, B, a*_0_, *τ*), column-wise *z*-scored across trials, and augmented with an intercept term, yielding **x**_*i*_ ∈ ℝ^*P*^ .

Because the response is constrained to (0, 1), we used a Beta regression with a logit link for the mean and a global concentration (precision) parameter. The model was

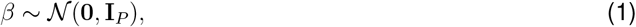

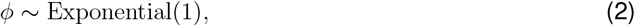

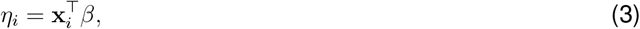

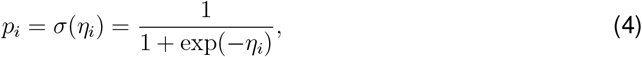

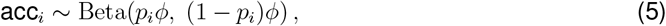

where *p*_*i*_ ∈ (0, 1) is the mean of the Beta likelihood and *ϕ >* 0 controls dispersion (larger *ϕ* implies lower variance around *p*_*i*_). For numerical stability, we bounded both observed accuracy values and the implied means away from 0 and 1 using a small constant *ϵ* = 10^−6^ (i.e., values were clamped to [*ϵ*, 1 − *ϵ*]).

We modeled trial-wise reaction times (RT; *rt*_*i*_ *>* 0) as a function of trial-wise drift diffusion model (DDM) parameter estimates. Let **x**_*i*_ ∈ ℝ^*P*^ denote the vector of standardized predictors for trial *i*, constructed from the DDM-derived covariates (*v, B, a*_0_, *τ*), where each column was *z*-scored across trials (mean 0, variance 1). An intercept term was appended so that 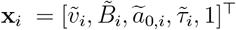 .

We used a Gamma generalized linear model with a log link to ensure the positivity of the mean RT. Specifically,

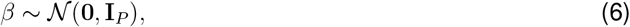

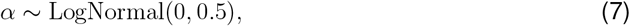

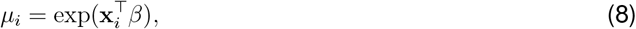

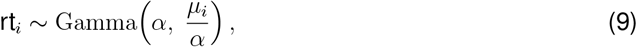

where *β* ∈ ℝ^*P*^ are the regression weights, *α >* 0 is the Gamma shape parameter, and we parameterize the Gamma likelihood by shape *α* and scale *µ*_*i*_*/α* so that E[rt_*i*_] = *µ*_*i*_.

For all models, we used the No-U-Turn-Sampler (NUTS) drawing 5000 samples from the posterior distribution. Posterior coefficient estimates and observed-versus-predicted accuracy and RT for these regressions are shown in **Supplemental Figure 7A–D**.

#### Model-based simulation checks

To evaluate model goodness-of-fit, we performed posterior predictive simulation using point estimates of model parameters. For each fitted model, we obtained the maximum a posteriori (MAP) estimate of the parameters and generated synthetic datasets under the generative model. Simulations preserved the empirical trial structure, including the number of trials per session. For the DDM–HMM, latent state sequences were first simulated from the inferred transition matrix, after which reaction times (RTs) and choices were generated from the state-specific drift diffusion process.

For each statistic of interest (e.g., marginal RT distributions for correct and incorrect trials, autocorrelation functions), we generated multiple simulated datasets and computed the corresponding summary statistic. Empirical statistics were compared to the simulated distribution, and credibility intervals were obtained by computing percentile bounds (typically 2.5%–97.5%) across simulations. Model adequacy was assessed by determining whether empirical values fell within these simulation-derived intervals. Posterior predictive checks of this kind appear throughout the figures: marginal RT distributions and Q–Q plots are shown in **Figure 3B–C**, the state-occupancy autocorrelation function is compared to model simulations in **Figure 4F**, reaction-time autocorrelations across models are shown in **Figure 6B**, short-timescale population autocorrelations are shown in **Figure 7D**, marginal RT distributions across all animals are shown in **Supplemental Figure 5**, and the same set of checks for an expert rat is shown in **Supplemental Figure 6C, G**.

#### State enrichment analyses

To determine whether specific behavioral conditions or task variables were overrepresented in particular latent states, we computed enrichment scores relative to chance expectations. For each state *s* and condition *c*, enrichment was defined as

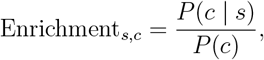

where *P* (*c* | *s*) is the empirical probability of condition *c* among trials assigned to state *s*, and *P* (*c*) is its marginal probability across all trials. Values greater than 1 indicate overrepresentation, and values less than 1 indicate depletion relative to chance. Enrichment of DDM-HMM state occupancy during the dark phase and during feeding is shown in **Figure 5B–C**.

#### Posterior alignment between models

To quantify similarity between latent states inferred by the GLM–HMM and the DDM–HMM, we compared their trial-wise posterior state probabilities. Let 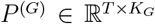 and 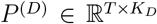 denote posterior matrices for the GLM–HMM and DDM–HMM, respectively, where rows correspond to trials and rows sum to 1.

We computed a soft co-occupancy (confusion) matrix

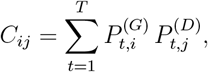

which represents the expected number of trials jointly assigned to GLM–HMM state *i* and DDM–HM state *j*. This approach avoids hard MAP discretization and preserves posterior uncertainty.

#### Chance-corrected alignment (lift)

Because marginal state occupancies differ across models, raw co-occupancy values are not directly interpretable. We therefore computed a chance-corrected enrichment (“lift”) matrix. Let *N* = Σ_*ij*_ *C*_*ij*_ and define row and column marginals *r*_*i*_ = Σ_*j*_ *C*_*ij*_ and *c*_*j*_ = Σ_*i*_ *C*_*ij*_. Under independence, the expected joint occupancy is

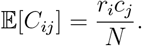

We defined lift as

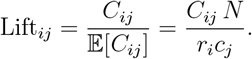

Values greater than 1 indicate above-chance alignment between the corresponding states. The lift matrix between GLM-HMM and DDM-HMM states is shown in **Supplemental Figure 11A**.

#### Normalized mutual information

To quantify alignment between trial-wise state assignments inferred by the DDM–HMM and behavioral labels derived from the GLM, we computed the normalized mutual information (NMI) between the two categorical sequences. Let *S* denote the discrete latent state inferred by the HMM and *G* the GLM-derived classification. Mutual information was defined as

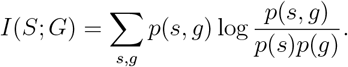

Normalization was performed using the symmetric formulation

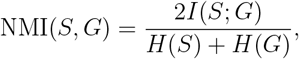

where *H*(·) denotes Shannon entropy. Probabilities were estimated empirically from trial frequencies. This normalization bounds the statistic between 0 (independent assignments) and 1 (perfect alignment).

To assess whether alignment exceeded chance, we generated null distributions by permuting trial indices of one model’s posterior matrix while preserving marginal occupancies. Empirical NMI values were compared to this null distribution. Multiple comparisons across animals were controlled using the Benjamini–Hochberg procedure with false discovery rate (FDR) set to 0.01. The NMI permutation test between GLM-HMM and DDM-HMM posteriors is shown in **Supplemental Figure 11B**.

### 7.9 Software

All analyses were conducted using Julia^46^ and R^41^.

Model development, inference, and simulation were implemented in Julia. The computational environment relied on the following core packages: Optim.jl^45^ (numerical optimization), ForwardDiff.jl^44^ (automatic differentiation), HiddenMarkovModels.jl^47^ (state-space inference), StateSpaceDynamics.jl^48^ (GLM-HMM learning and inference), Turing.jl^49^ (probabilistic modeling), and Plots.jl^50^ and Makie.jl^51^ (visualization). These libraries were used for parameter estimation via Expectation–Maximization, likelihood evaluation, gradient-based optimization, posterior inference, simulation from fitted models, and generation of figures.

Statistical analyses of behavioral modulation across the 24-hour cycle were performed in R using mgcv^40^ for generalized additive mixed modeling (GAMM) and ggplot2^52^ for visualization.

## 8 RESOURCE AVAILABILITY

### 8.1 Lead contact

Requests for further information and resources should be directed to and will be fulfilled by the lead contact, BBS and BDD.

### 8.2 Materials availability

This study did not generate new materials.

### 8.3 Data and code availability

All custom code used for data processing, model fitting, simulation, and figure generation is publicly available. The full analysis pipeline and behavioral dataset are available at:

- https://github.com/depasquale-lab/24_Hour_Behavior
- https://github.com/depasquale-lab/DriftDiffusionModels.jl

The `24_Hour_Behavior` repository contains the complete dataset required to reproduce all results reported in this study, along with scripts for preprocessing, model fitting, and figure generation. Analyses necessary for reproducing the results are included in the repository as plain Julia scripts, R scripts, and Pluto notebooks.^53^ The `DriftDiffusionModels.jl` repository contains the implementation of the DDM and DDM-HMM frameworks used in this work.

## 9 ACKNOWLEDGMENTS

We thank Christa Rose for help with daily rat training. We also thank Chandramouli Chandrasekaran and Grant McConachie for reading and providing valuable comments on the manuscrip This work was supported by NIMH award number R56MH132732 to BBS, a Scialog Grant from the Research Corporation for Science Advancement to BD, a Center for Systems Neuroscience distinguished fellowship award to GAK, a Fulbright/FLAD Research Award to HFD, a student research fellowship (UROP) from Boston University to HFD and JF, a Hariri Graduate Student Fellowship to RAS, and a Ludwig Family Foundation grant to SR.

## 10 AUTHOR CONTRIBUTIONS

HX, QD, GAK, and BBS designed the experiments. HX, HFD, QD performed the experiments. RAS and BD designed the model-based analyses with input from BBS. RAS, HX, and HFD performed analysis. BBS, BDD, and SR provided research support. RAS wrote the manuscript with input from HFD, SR, BD, and BBS. All authors approved the final manuscript.

## 13 Supplemental Figures

**Supplemental Figure 1:**
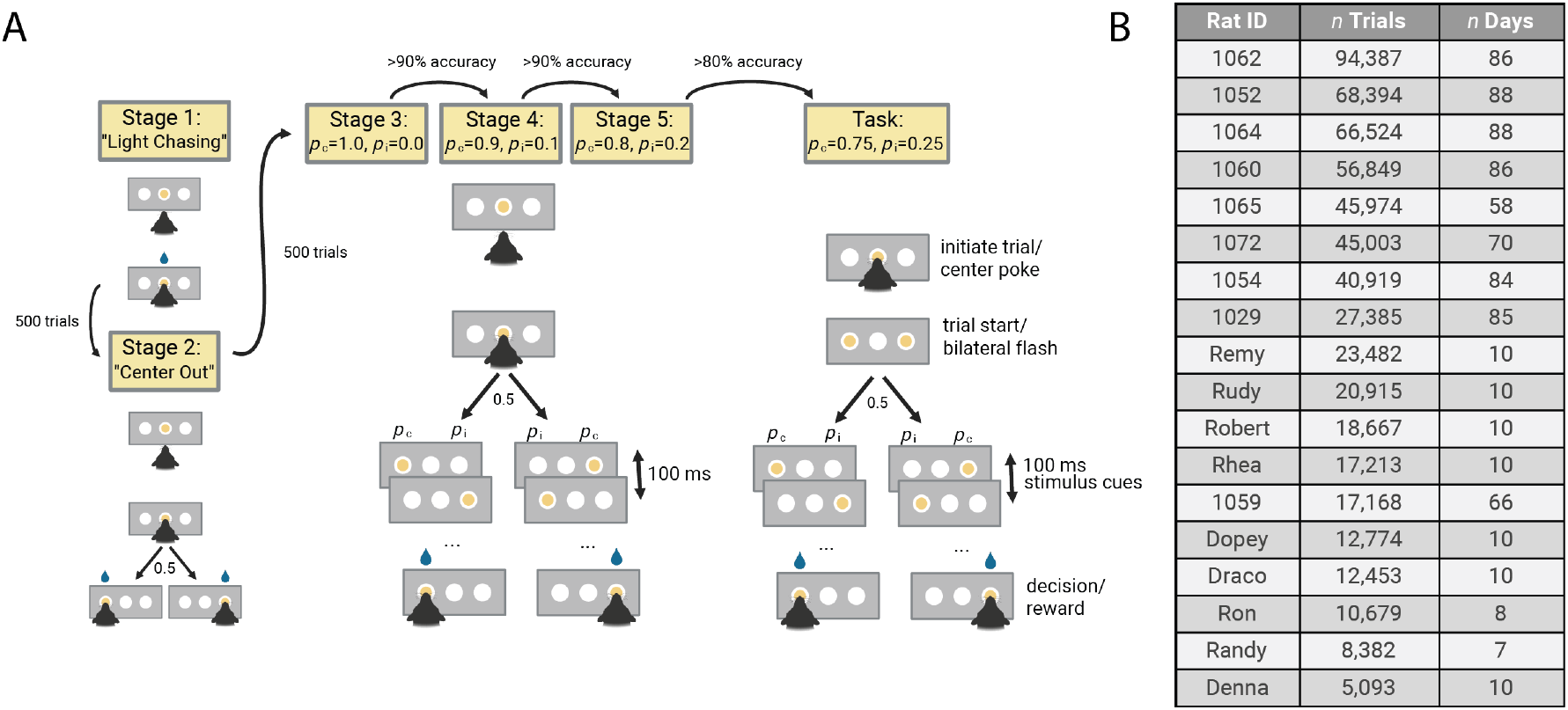
Animal training procedure and dataset summary. **(A)** Animals are trained in a six-stage procedure. In stage 1 (light chasing), animals nose poke the illuminated center port to receive a reward (500 trials). In stage 2 (center-out), animals nose poke the center port and then either the left or right port (assigned randomly; 500 trials). In stages 3–5, animals initiate trials via the center port and must achieve *>* 90% accuracy over 500 trials by selecting the port with the higher flash probability. Before advancing to the performance stage (stage 6), animals must achieve *>* 80% accuracy over 500 trials. **(B)** Summary statistics for the 18 rats included in the study.

**Supplemental Figure 2:**
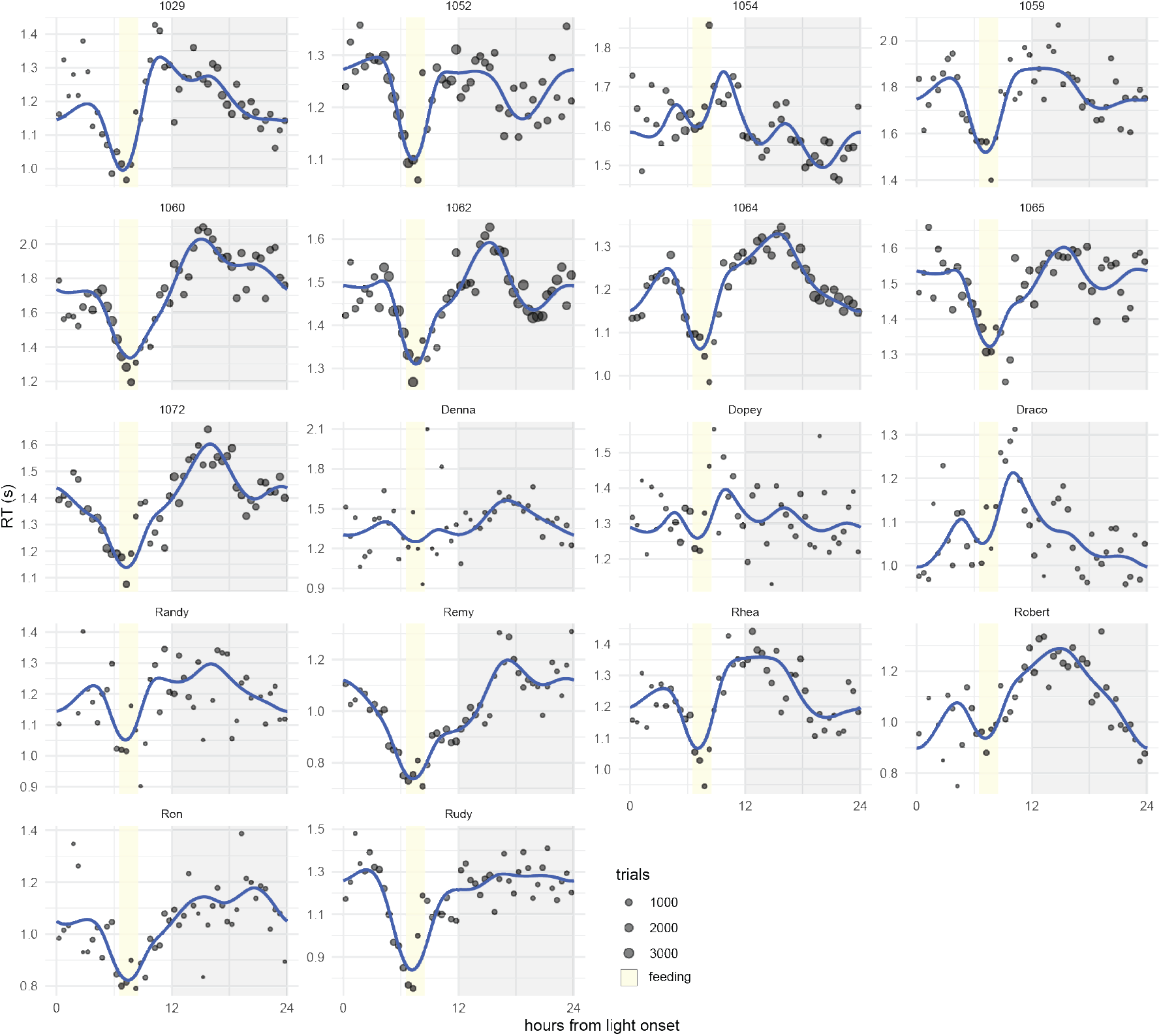
Individual GAM fits of reaction times across the 24-hour cycle. Blue lines indicate animal-level averages; points are scaled by the number of trials contributing to each estimate; yellow bars denote feeding periods.

**Supplemental Figure 3:**
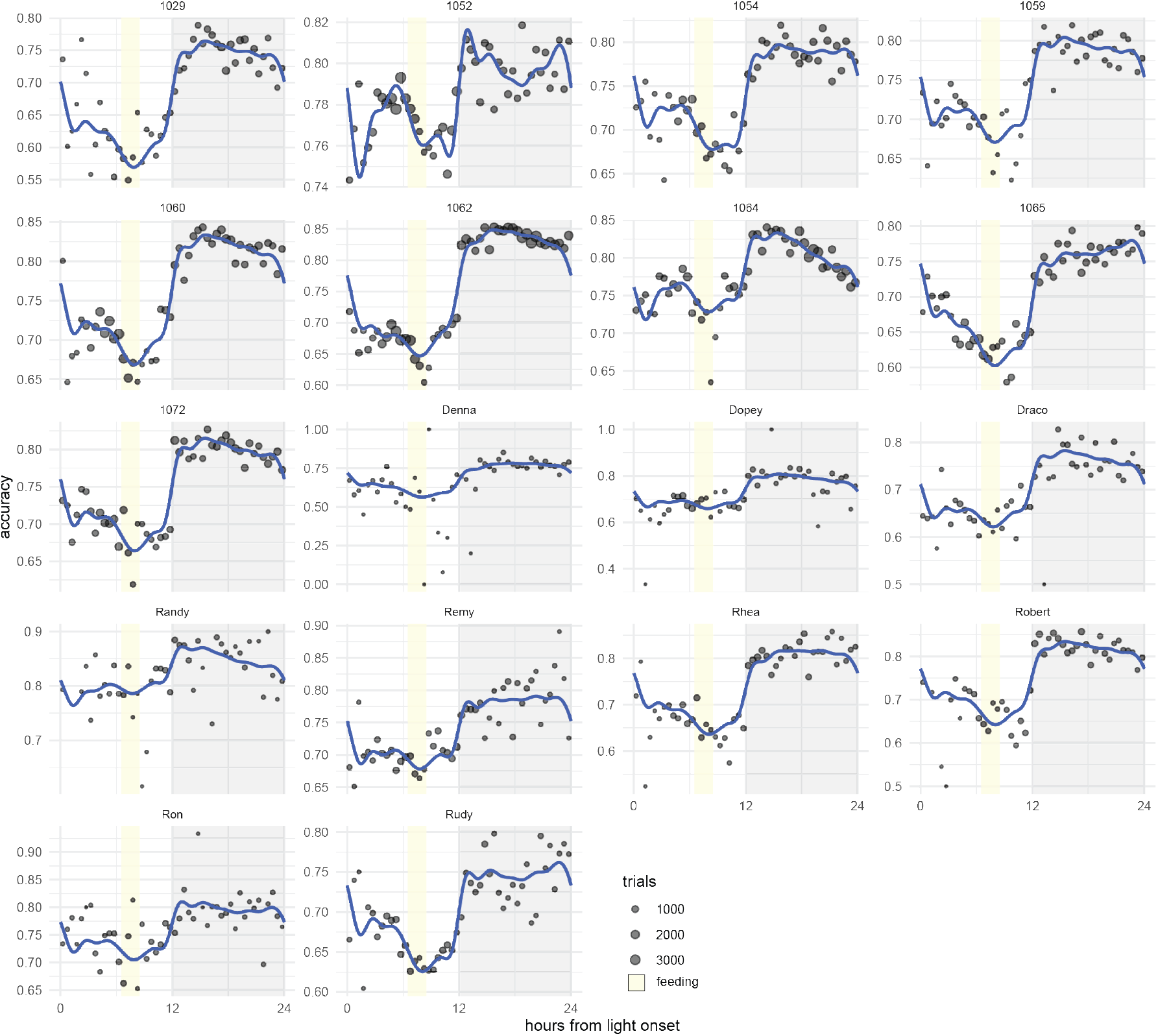
Individual GAM fits of accuracy across the 24-hour cycle. Blue lines indicate animal-level averages; points are scaled by the number of trials; yellow bars denote feeding periods.

**Supplemental Figure 4:**
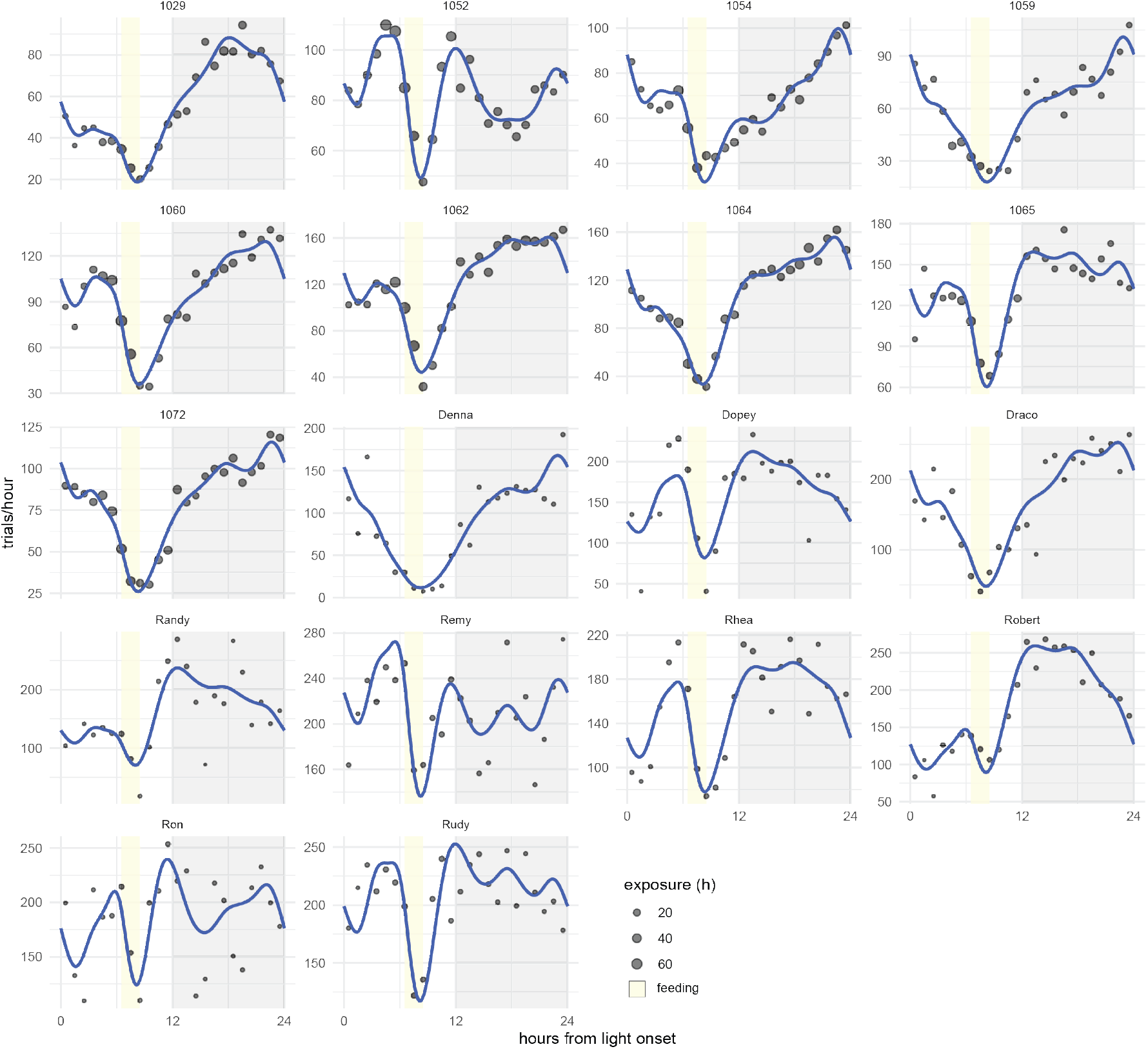
Individual GAM fits of trial production across the 24-hour cycle. Blue lines indicate animal-level averages; points are scaled by the number of trials; yellow bars denote feeding periods.

**Supplemental Figure 5:**
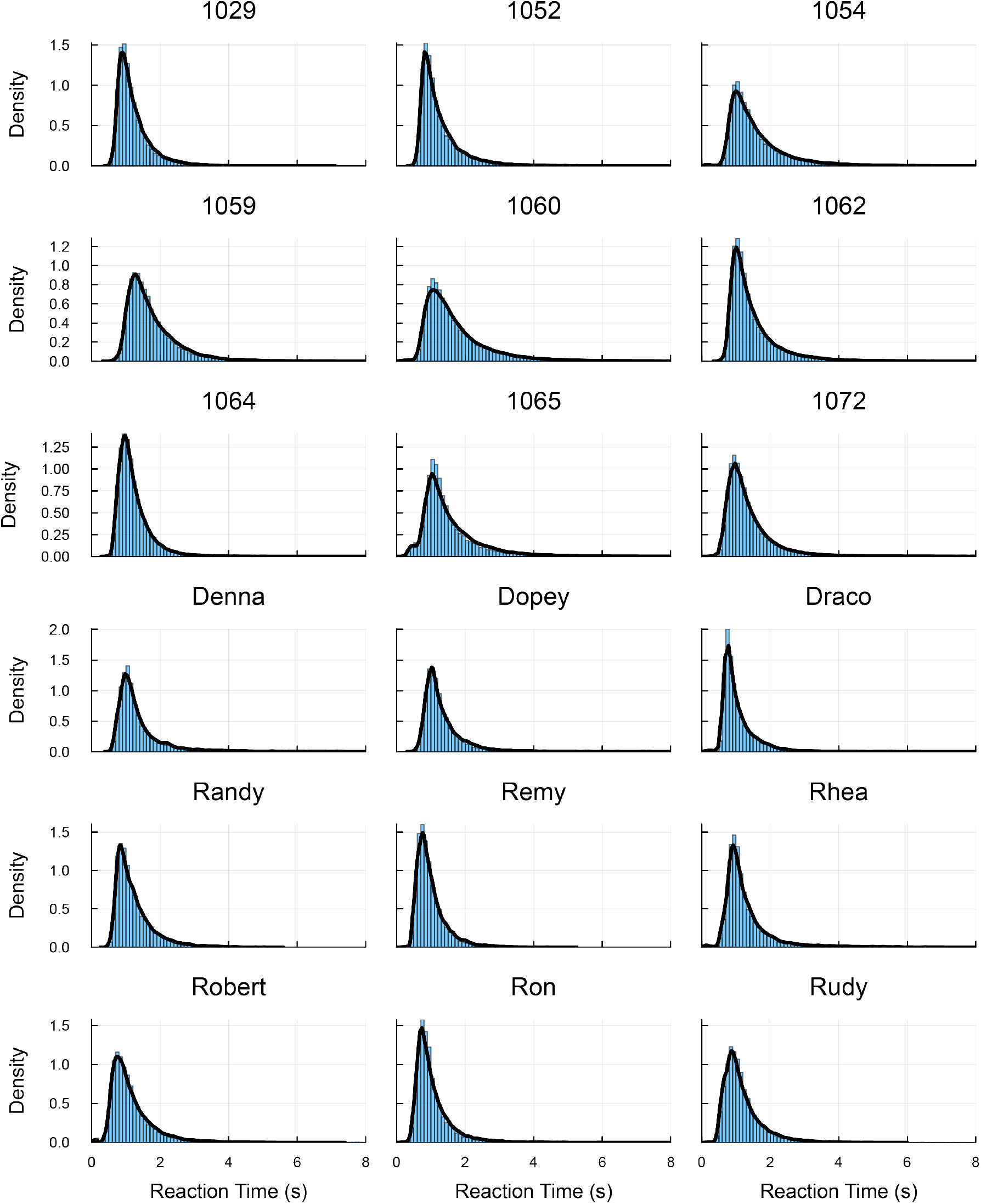
The DDM-HMM captures RT distributions. Simulations from learned DDM-HMMs (black line) closely match the empirical data (blue histogram) from all animals.

**Supplemental Figure 6:**
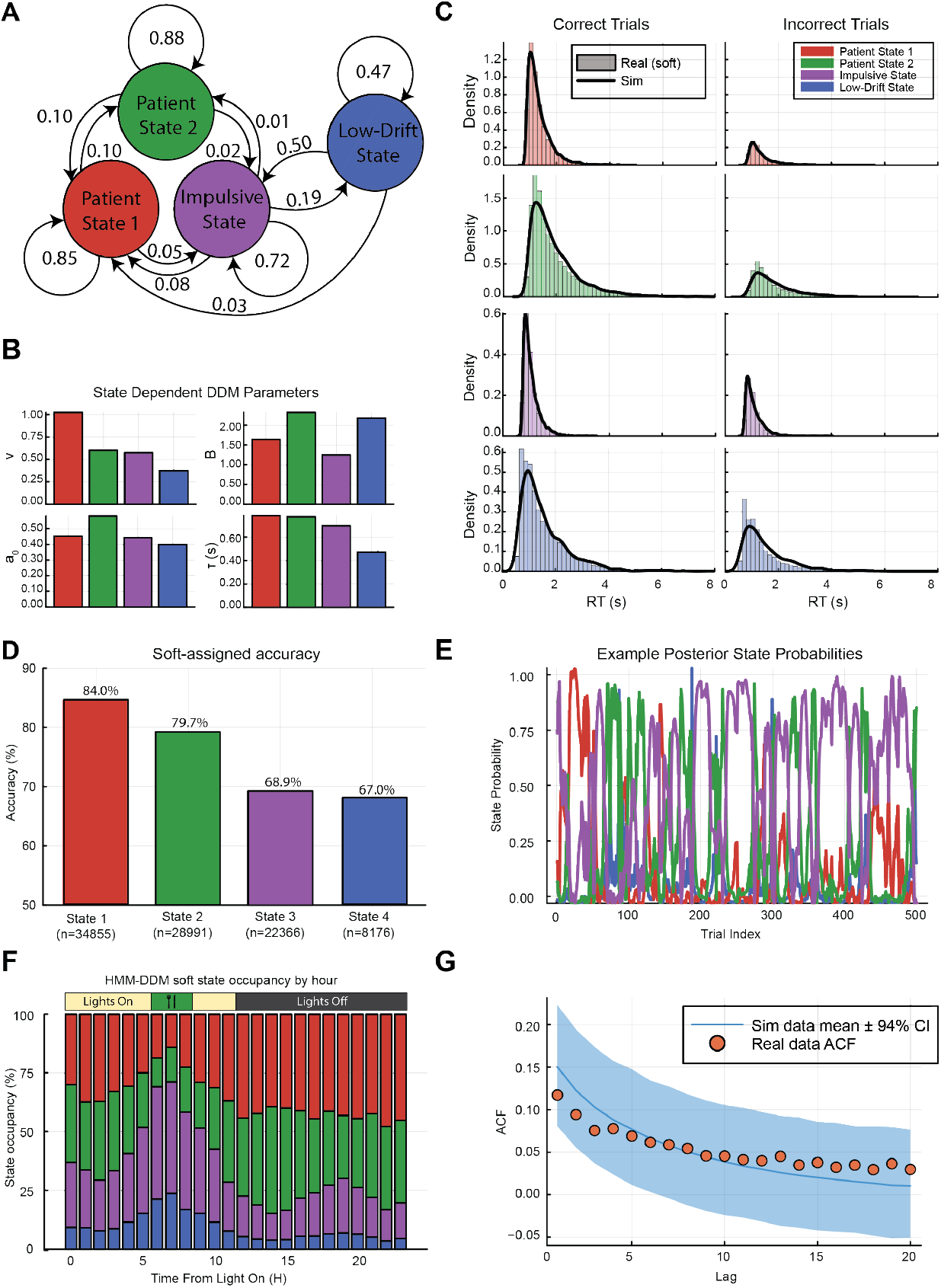
Example DDM-HMM fit for an expert rat. **(A)** Schematic of the transition dynamics between the four inferred latent decision states: patient state 1, patient state 2, impulsive state, and low-drift. **(B)** State-dependent drift–diffusion model (DDM) parameters (drift rate, boundary separation, starting bias, and non-decision time), illustrating systematic differences in decision dynamics across latent states. **(C)** Reaction time (RT) distributions for correct/incorrect trials, shown for each state. Histograms reflect real data using soft state assignments; black curves indicate simulated data from the fitted HMM–DDM. **(D)** Soft-assigned choice accuracy for each state, demonstrating higher accuracy in patient states and reduced accuracy in impulsive and low-drift states. **(E)** Example posterior state probabilities across trials, showing rapid switching and extended epochs of state dominance. **(F)** Hourly state occupancy relative to the light–dark cycle, revealing circadian modulation of latent decision states. **(G)** Autocorrelation function (ACF) of inferred states in the real data (points) compared with simulated data from the model (mean *±* 94% CI), indicating that the model captures temporal structure in state transitions.

**Supplemental Figure 7:**
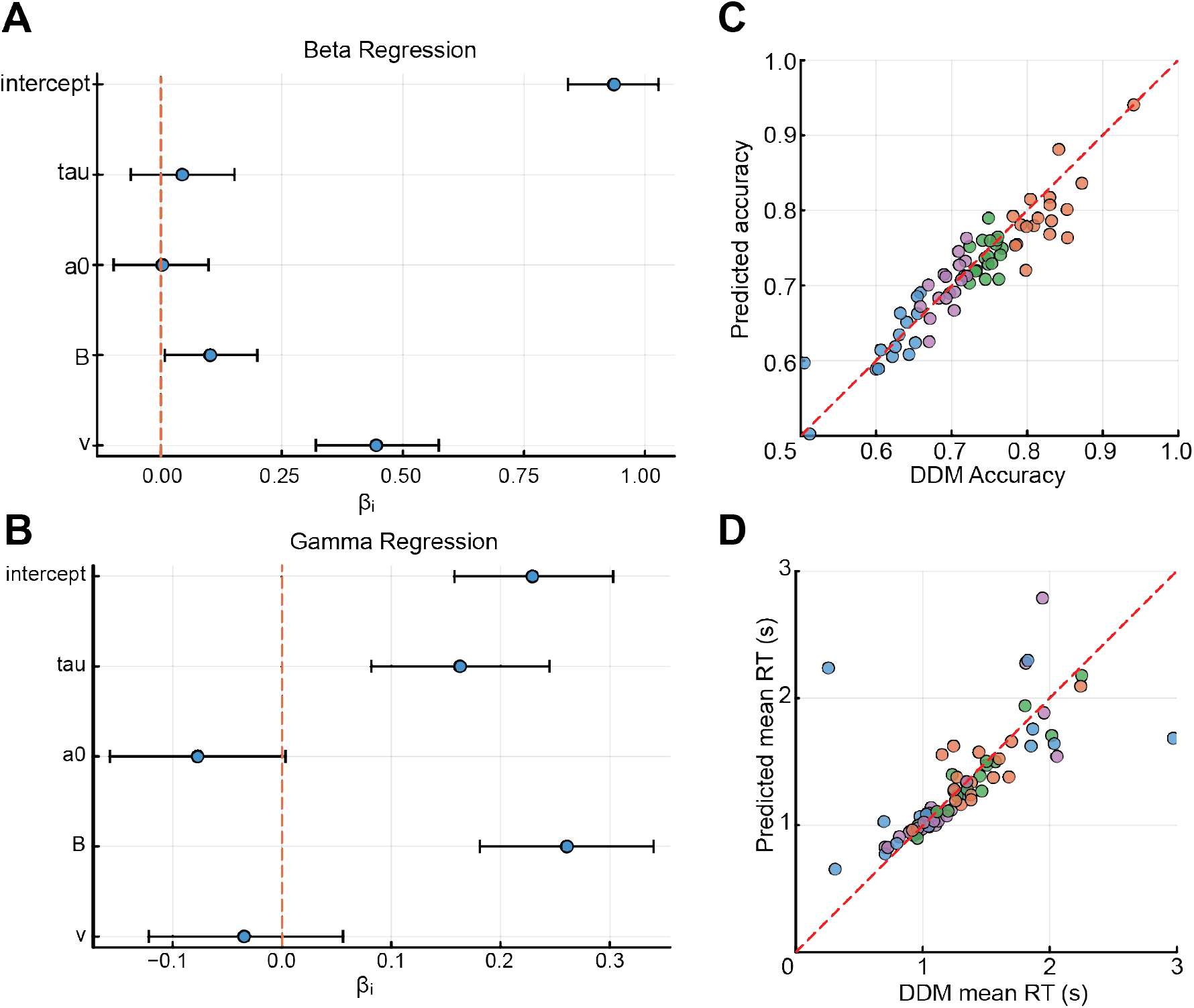
DDM parameter regressions. **(A)** Forest plot of posterior mean coefficients and 95% credible intervals for beta regression predicting accuracy from DDM parameters. **(B)** Forest plot of posterior mean coefficients and 95% credible intervals predicting mean reaction time from DDM parameters. **(C)** Observed versus predicted accuracy from the beta regression. **(D)** Observed versus predicted reaction time from the gamma regression.

**Supplemental Figure 8:**
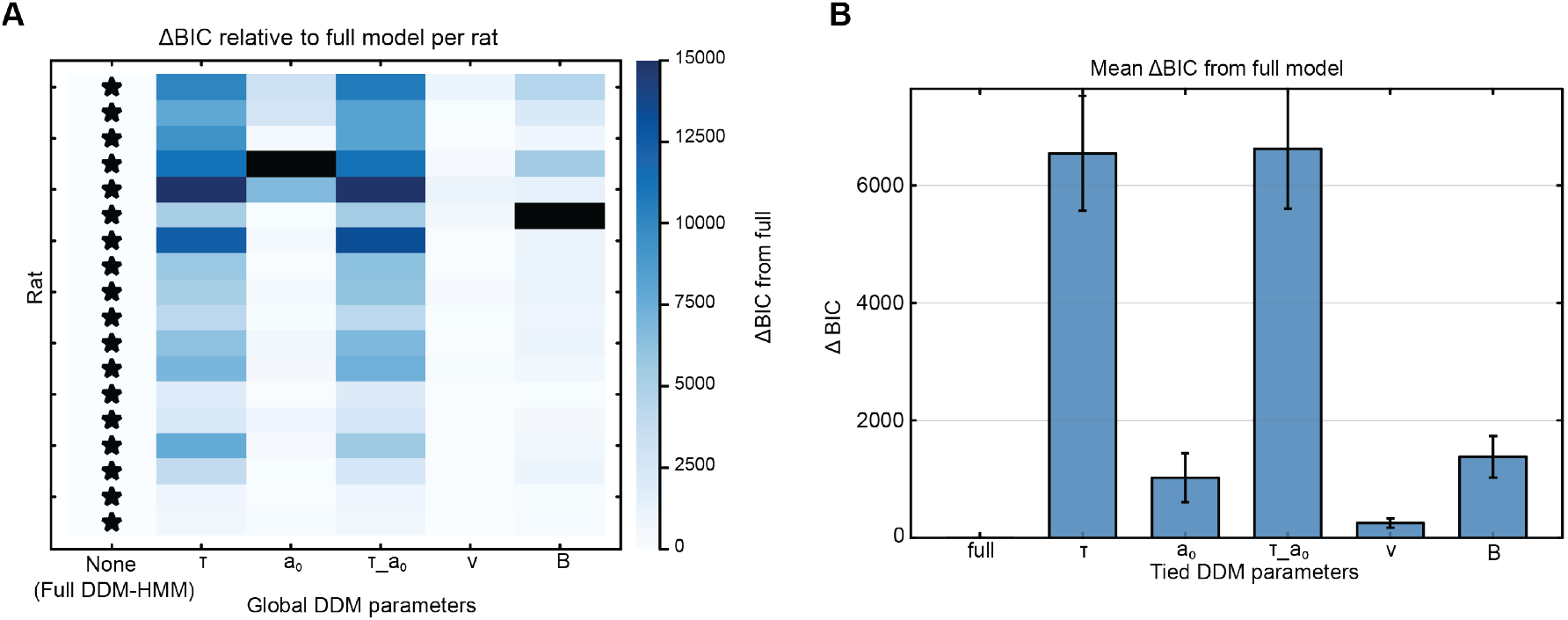
BIC comparison suggests the full DDM-HMM is the best model of animal 24-hour decision making data. **(A)** Heatmap of individual animal ΔBIC from the “full” model. Black stars indicate the best fitting model. Bluer colors represent worse model fits compared to the full model. All individual animal data is best fit by the full DDM-HMM. X-axis indicates which model parameters were fit globally i.e., shared across states. **(B)** Mean ΔBIC at the population level. All other candidate models are worse than the full DDM-HMM. X-axis indicates which model parameters were fit globally i.e., shared across states.

**Supplemental Figure 9:**
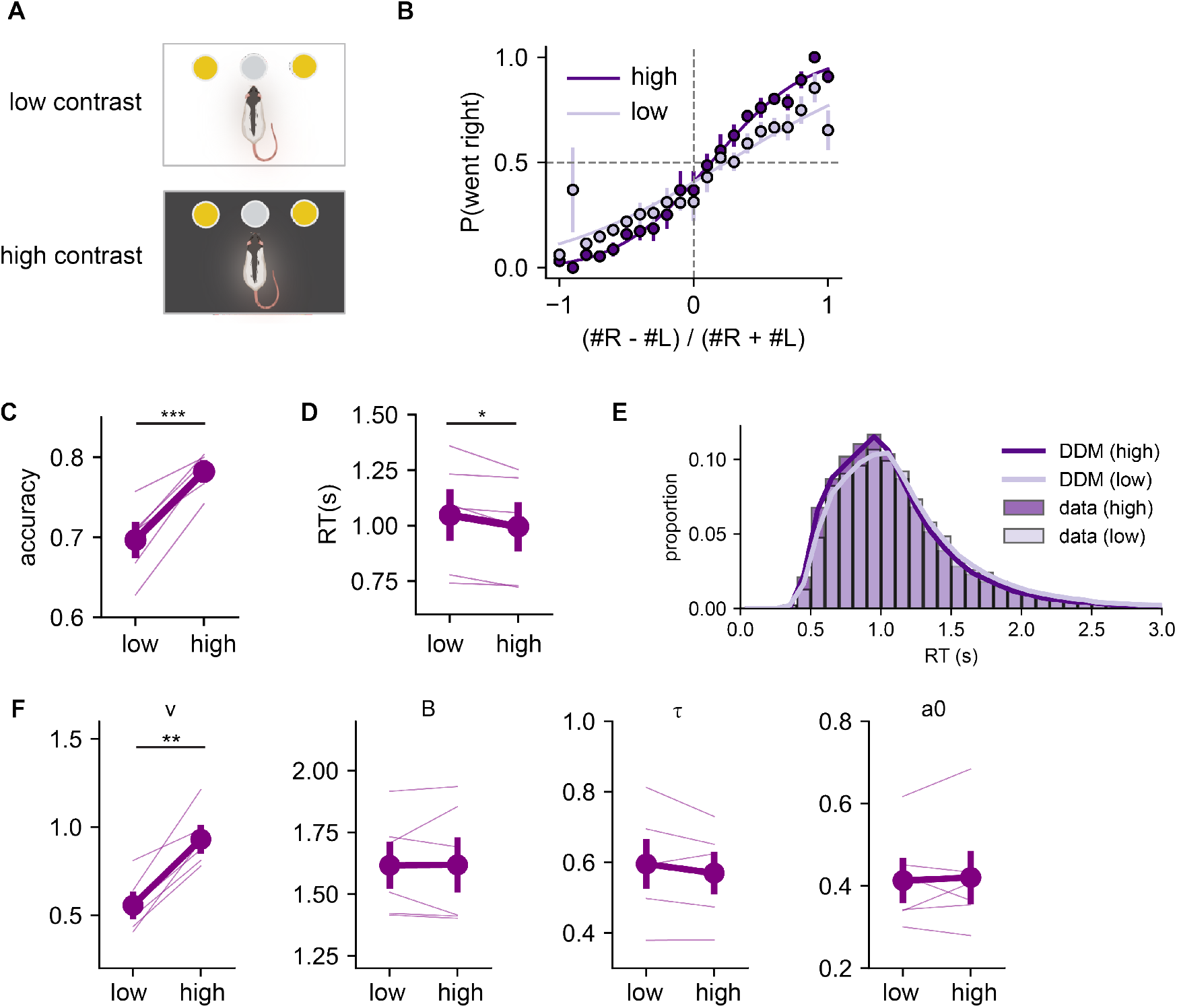
Rats show a higher drift rate under high contrast conditions during daily 2-hr training. **(A)** Schematic of rats training under low contrast (room light on) and high contrast (room light off). **(B)** Psychometric curves of rats performing the task under low and high contrast conditions. **(C)** Average accuracy across animals (*±* SE; *n* = 6). Each line represents the mean accuracy from an individual animal. **(D)** Average RT across animals (*±* SE; *n* = 6). Each line represents median RT across animal. **(E)** RT distributions from low and high contrast. Bars represent the data, lines represent DDM predictions. **(F)** Mean fitted DDM parameters for drift rate (*v*), boundary separation (*B*), non-decision time (*τ*) and starting point (*a*_0_).

**Supplemental Figure 10:**
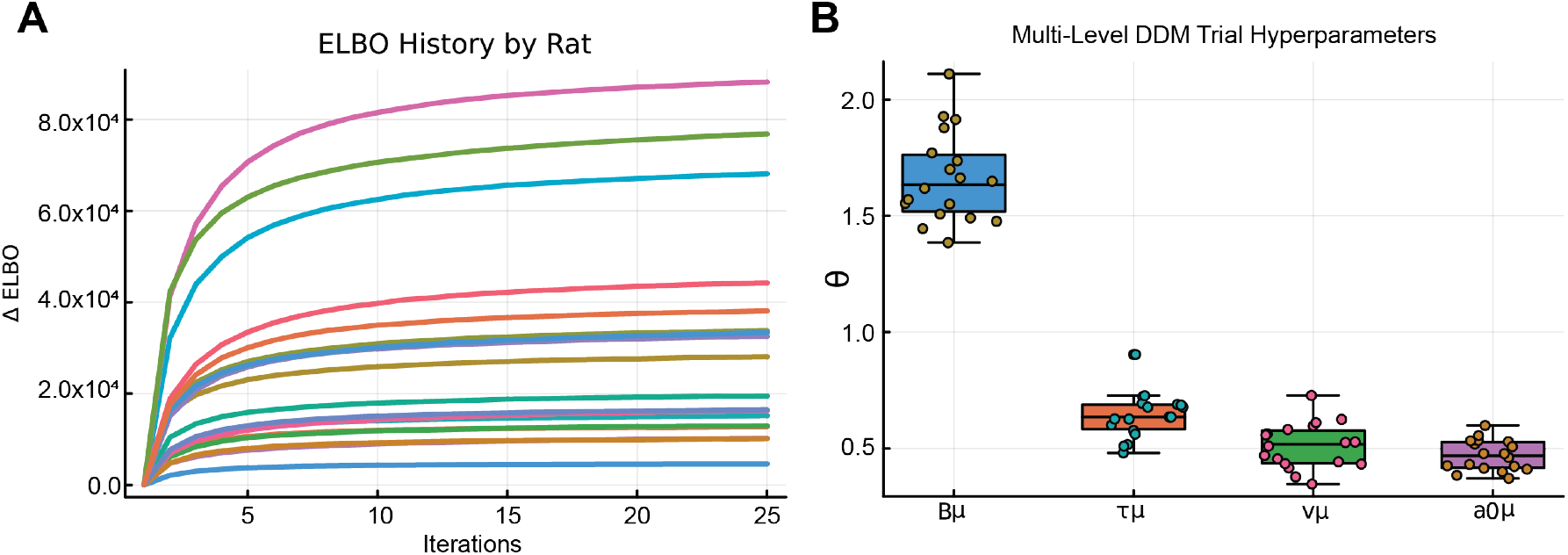
Parameter learning in a multi-level DDM model. **(A)** Change in evidence lower bound (ΔELBO) across optimization iterations for each rat, demonstrating stable convergence of the multi-level DDM fits. Each colored trace corresponds to a single animal. **(B)** Posterior distributions of trial-level hyperparameters from the multilevel DDM, including boundary height (*β*_*µ*_), non-decision time (*τ*_*µ*_), drift rate (*v*_*µ*_), and starting-point bias (*a*_0*µ*_), pooled across animals.

**Supplemental Figure 11:**
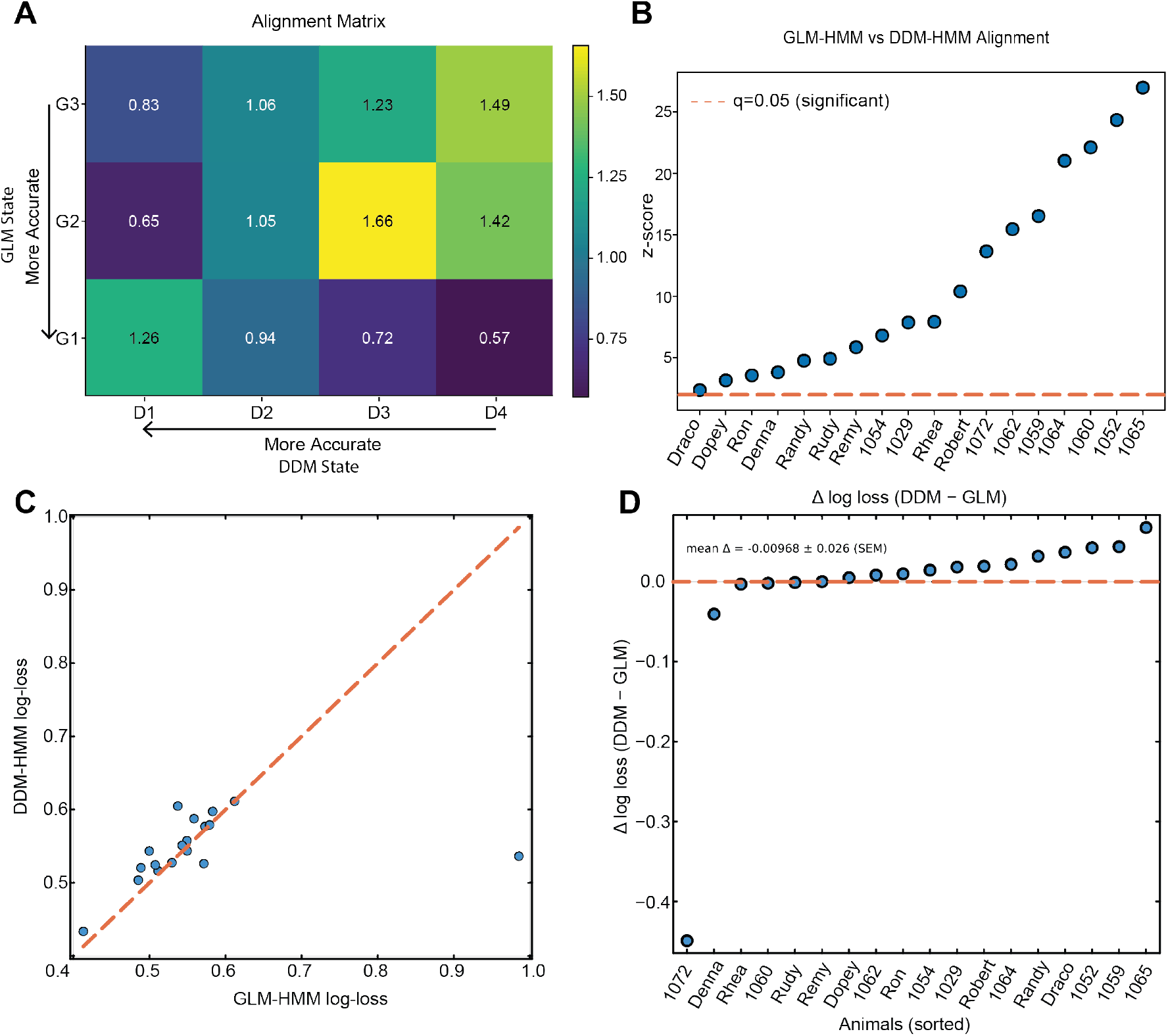
The DDM-HMM and GLM-HMM find similar latent dynamics. **(A)** Alignment matrix showing the how many more times above chance two states are jointly active. **(B)** Permutation test results calculating NMI between the GLM-HMM and DDM-HMM posteriors. All but three animals display significant mutual information. **(C)** When using state posteriors to calculate the probability of correct choices, both models have similar predictive performance. **(D)** Quantification of results in **C** i.e., the difference between the log-loss for both models. The distribution is centered at zero indicating no superior model fit.

**Supplemental Figure 12:**
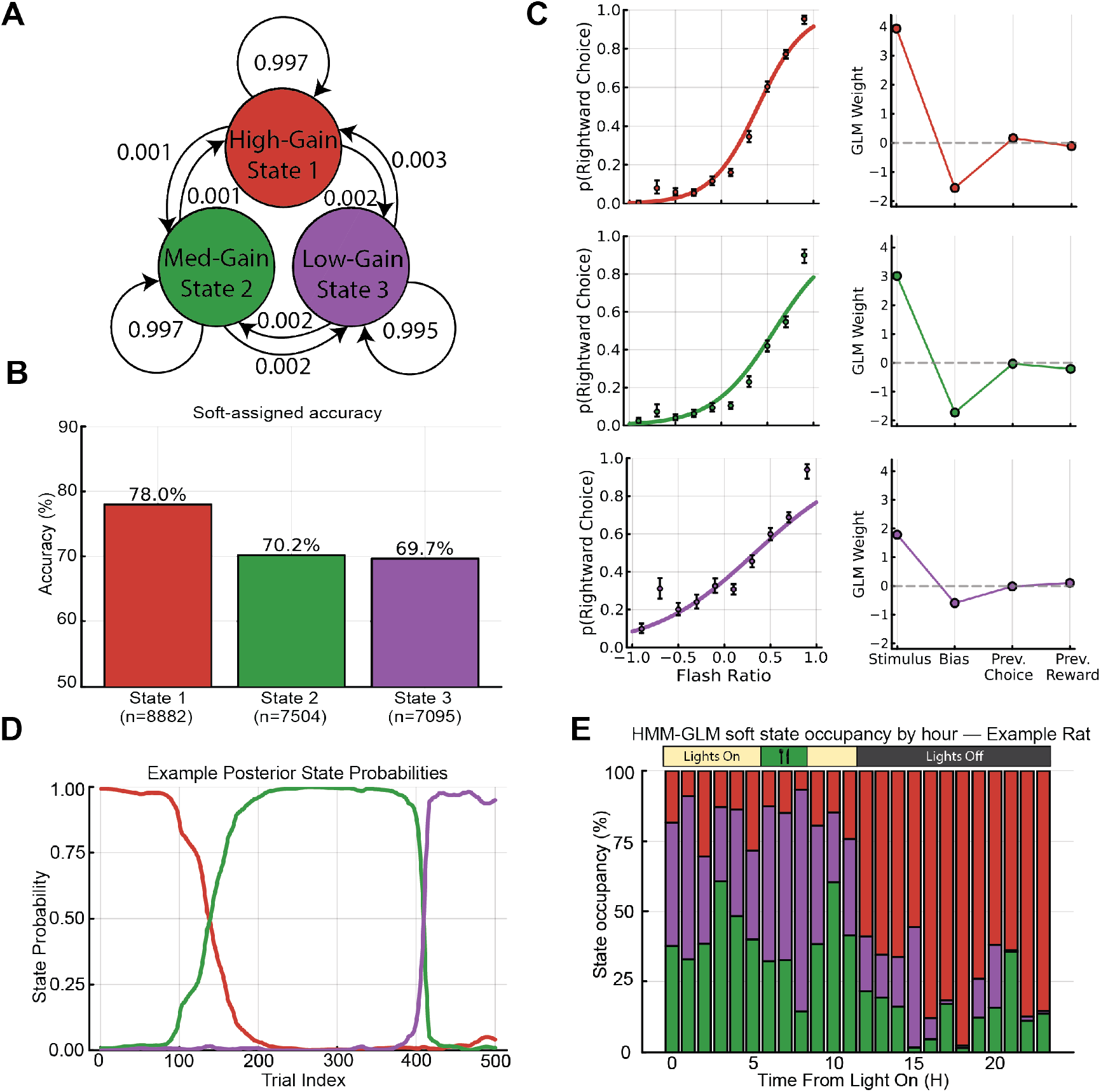
GLM-HMM latent states align with task structure. **(A)** State transition diagram for a three-state GLM-HMM fit to an example rat. Nodes indicate latent states, colored by inferred gain (sensitivity to stimulus evidence), and arrows denote transition probabilities between states. **(B)** Soft-assigned choice accuracy for each GLM-HMM state, demonstrating a graded hierarchy of performance across states. **(C)** Psychometric functions (left) and corresponding GLM weights (right) for each state, showing systematic differences in stimulus sensitivity and history dependence across latent states. **(D)** Example posterior state probabilities across successive trials, illustrating strong state persistence and discrete switching behavior. **(E)** State occupancy as a function of time from lights on, revealing systematic modulation of GLM-HMM states across the circadian cycle.

**Supplemental Figure 13:**
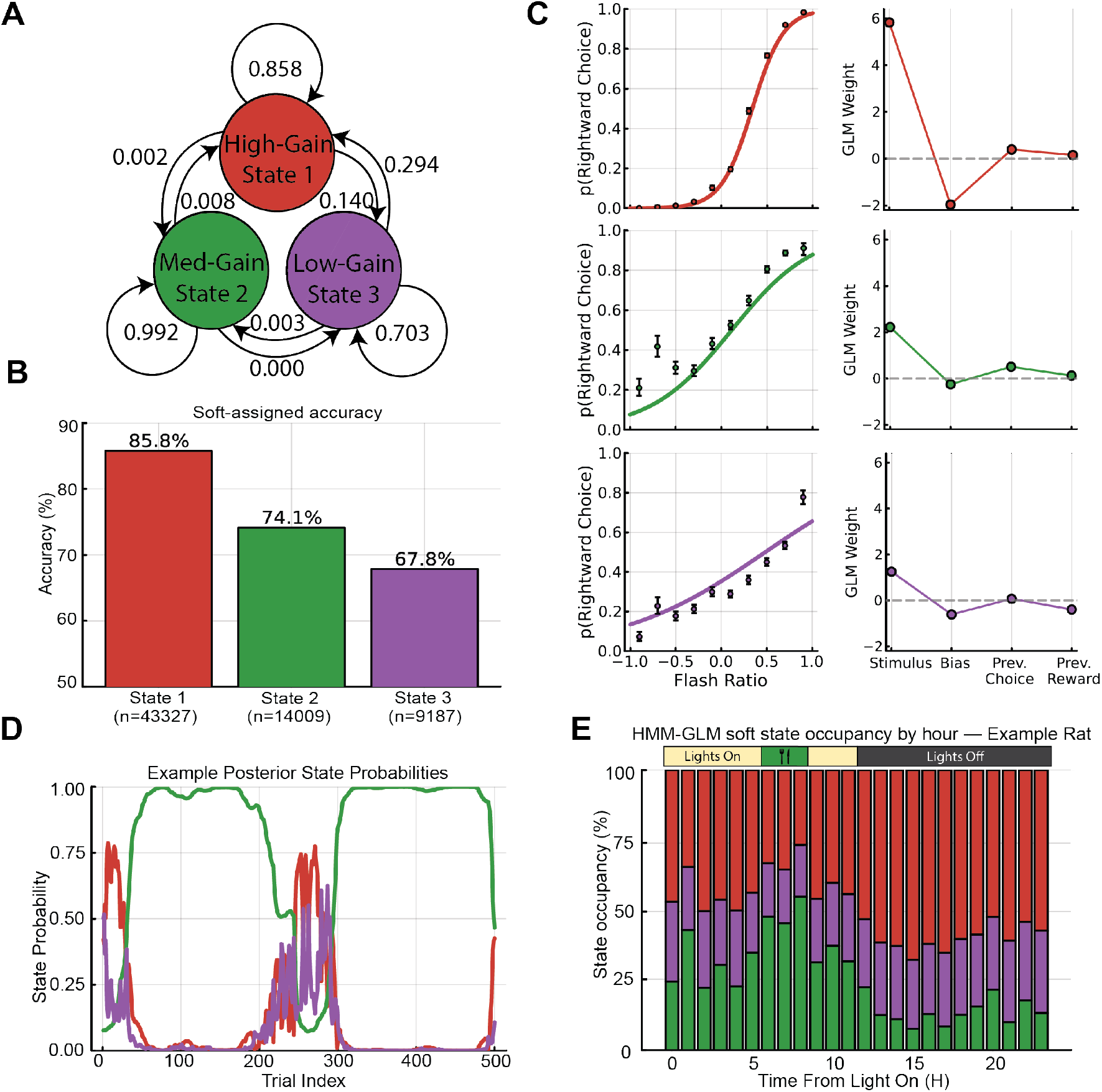
Replication of GLM-HMM state structure in an expert rat. **(A)** State transition structure for a three-state GLM-HMM, showing high self-transition probabilities indicative of persistent latent states. **(B)** Soft-assigned accuracy across states, again revealing a graded performance hierarchy. **(C)** State-specific psychometric functions and GLM weights, demonstrating consistent modulation of stimulus gain and weak history effects across states. **(D)** Posterior state probabilities across trials, showing extended persistence within states. **(E)** Circadian modulation of GLM-HMM state occupancy, with higher-gain states predominating during the dark phase and lower-gain states enriched during feeding and light periods.

## References

1. Rothschild, J.M., Keohane, C.A., Rogers, S., Gardner, R., Lipsitz, S.R., Salzberg, C.A., Yu, T., Yoon, C.S., Williams, D.H., Wien, M.F., Czeisler, C.A., Bates, D.W., and Landrigan, C.P. (2009). Risks of complications by attending physicians after performing nighttime procedures. JAMA 302, 1565–1572. doi: 10.1001/jama.2009.1423.

2. Landrigan, C.P. (2010). Effect of lack of sleep on medical errors. In F.P. Cappuccio, M.A. Miller, and S.W. Lockley, eds. Sleep, Health and Society: From Aetiology to Public Health chap. 17. pp. 382–396. Oxford University Press. ISBN 9780199566594 pp. 382–396. doi: 10.1093/acprof:oso/9780199566594.003.0017.

3. Lenné, M.G., Triggs, T.J., and Redman, J.R. (1997). Time of day variations in driving performance. Accident Analysis & Prevention 29, 431–437. doi: 10.1016/S0001-4575(97)00022-5.

4. International Brain Laboratory, Angelaki, D., Benson, B., Benson, J., Birman, D., Bonacchi, N., Bougrova, K., Bruijns, S.A., Carandini, M., Catarino, J.A., Chapuis, G.A., Churchland, A.K., Dan, Y., Davatolhagh, F., Dayan, P., DeWitt, E.E., Engel, T.A., Fabbri, M., Faulkner, M., Fiete, I.R., Findling, C., Freitas-Silva, L., Gerçek, B., Harris, K.D., Häusser, M., Hofer, S.B., Hu, F., Hubert, F., Huntenburg, J.M., Khanal, A., Krasniak, C.S., Langdon, C., Langfield, C., Lau, P.Y.P., Mainen, Z.F., Meijer, G.T., Miska, N.J., Mrsic-Flogel, T.D., Noel, J.P., Nylund, K., Pan-Vazquez, A., Paninski, L., Pouget, A., Rossant, C., Roth, N., Schaeffer, R., Schartner, M., Shi, Y., Socha, K.Z., Steinmetz, N.A., Svoboda, K., Urai, A.E., Wells, M.J., West, S.J., Whiteway, M.R., Winter, O., and Witten, I.B. (2025). A brain-wide map of neural activity during complex behaviour. Nature 645, 177–191. doi: 10.1038/s41586-025-09235-0.

5. Findling, C., Hubert, F., International Brain Laboratory, Acerbi, L., Benson, B., Benson, J., Birman, D., Bonacchi, N., Buchanan, E.K., Bruijns, S., Carandini, M., Catarino, J.A., Chapuis, G.A., Churchland, A.K., Dan, Y., Davatolhagh, F., DeWitt, E.E.J., Engel, T.A., Fabbri, M., Faulkner, M.A., Fiete, I.R., Freitas-Silva, L., Gerçek, B., Harris, K.D., Häusser, M., Hofer, S.B., Hu, F., Huntenburg, J.M., Khanal, A., Krasniak, C., Langdon, C., Langfield, C.A., Latham, P.E., Lau, P.Y.P., Mainen, Z., Meijer, G.T., Miska, N.J., Mrsic-Flogel, T.D., Noel, J.P., Nylund, K., Pan-Vazquez, A., Paninski, L., Pillow, J., Rossant, C., Roth, N., Schaeffer, R., Schartner, M., Shi, Y., Socha, K.Z., Steinmetz, N.A., Svoboda, K., Tessereau, C., Urai, A.E., Wells, M.J., West, S.J., Whiteway, M.R., Winter, O., Witten, I.B., Zador, A., Zhang, Y., Dayan, P., and Pouget, A. (2025). Brain-wide representations of prior information in mouse decision-making. Nature 645, 192–200. doi: 10.1038/s41586-025-09226-1.

6. Luo, T.Z., Kim, T.D., Gupta, D., Bondy, A.G., Kopec, C.D., Elliott, V.A., DePasquale, B., and Brody, C.D. (2025). Transitions in dynamical regime and neural mode during perceptual decisions. Nature 646, 1156–1166. doi: 10.1038/s41586-025-09528-4.

7. Schiereck, S.S., Pérez-Rivera, D.T., Mah, A., DeMaegd, M.L., Hocker, D., Ward, R.M., Savin, C., and Constantinople, C.M. (2026). The orbitofrontal cortex updates beliefs for state inference. Neuron 114, 507–520.e8. doi: 10.1016/j.neuron.2025.10.024.

8. Finkelstein, A., Daie, K., Rózsa, M., Darshan, R., and Svoboda, K. (2026). Connectivity underlying motor cortex activity during goal-directed behaviour. Nature 649, 416–422. doi: 10.1038/s41586-025-09758-6.

9. Maheu, M., and Scott, B.B. (2025). Guiding principles for shaping instructed behaviors in lab rodents. PsyArXiv. doi: 10.31234/osf.io/asdpu_v1.

10. Gritton, H.J., Sutton, B.C., Martinez, V., Sarter, M., and Lee, T.M. (2009). Interactions between cognition and circadian rhythms: attentional demands modify circadian entrainment. Behav. Neurosci. 123, 937–948. doi: 10.1037/a0017128.

11. Dijk, D.J., and von Schantz, M. (2005). Timing and consolidation of human sleep, wakefulness, and performance by a symphony of oscillators. J. Biol. Rhythms 20, 279–290. doi: 10.1177/0748730405278292.

12. Stephan, F.K. (1986). The role of period and phase in interactions between feeding- and light-entrainable circadian rhythms. Physiol. Behav. 36, 151–158. doi: 10.1016/0031-9384(86)90089-2.

13. Benstaali, C., Mailloux, A., Bogdan, A., Auzéby, A., and Touitou, Y. (2001). Circadian rhythms of body temperature and motor activity in rodents their relationships with the light-dark cycle. Life Sci. 68, 2645–2656. doi: 10.1016/S0024-3205(01)01081-5.

14. Francis, N.A., Bohlke, K., and Kanold, P.O. (2019). Automated behavioral experiments in mice reveal periodic cycles of task engagement within circadian rhythms. eNeuro 6, ENEURO.0121–19.2019. doi: 10.1523/ENEUR0.0121-19.2019.

15. Wyatt, J.K., Cecco, A.R.D., Czeisler, C.A., and Dijk, D.J. (1999). Circadian temperature and melatonin rhythms, sleep, and neurobehavioral function in humans living on a 20-h day. Am. J. Physiol. Regul. Integr. Comp. Physiol. 277, R1152–R1163. doi: 10.1152/ajpregu.1999.277.4.R1152.

16. Wyatt, J.K., Cajochen, C., Ritz-De Cecco, A., Czeisler, C.A., and Dijk, D.J. (2004). Low-dose repeated caffeine administration for circadian-phase-dependent performance degradation during extended wakefulness. Sleep 27, 374–381. doi: 10.1093/sleep/27.3.374.

17. Stephan, F.K. (1983). Circadian rhythms in the rat: constant darkness, entrainment to T cycles and to skeleton photoperiods. Physiol. Behav. 30, 451–462. doi: 10.1016/0031-9384(83)90152-X.

18. Cuesta, M., Clesse, D., Pévet, P., and Challet, E. (2009). From daily behavior to hormonal and neurotransmitters rhythms: comparison between diurnal and nocturnal rat species. Horm. Behav. 55, 338–347. doi: 10.1016/j.yhbeh.2008.10.015.

19. Bogacz, R., Brown, E., Moehlis, J., Holmes, P., and Cohen, J.D. (2006). The physics of optimal decision making: a formal analysis of models of performance in two-alternative forced-choice tasks. Psychol. Rev. 113, 700–765. doi: 10.1037/0033-295X.113.4.700.

20. Ratcliff, R., and McKoon, G. (2008). The diffusion decision model: theory and data for two-choice decision tasks. Neural Comput. 20, 873–922. doi: 10.1162/neco.2008.12-06-420.

21. Calhoun, A.J., Pillow, J.W., and Murthy, M. (2019). Unsupervised identification of the internal states that shape natural behavior. Nat. Neurosci. 22, 2040–2049. doi: 10.1038/s41593-019-0512-x.

22. Roy, N.A., Bak, J.H., Akrami, A., Brody, C.D., and Pillow, J.W. (2021). Extracting the dynamics of behavior in sensory decision-making experiments. Neuron 109, 597–610.e6. doi: 10.1016/j.neuron.2020.12.004.

23. Bolkan, S.S., Stone, I.R., Pinto, L., Ashwood, Z.C., Iravedra Garcia, J.M., Herman, A.L., Singh, P., Bandi, A., Cox, J., Zimmerman, C.A., Cho, J.R., Engelhard, B., Pillow, J.W., and Witten, I.B. (2022). Opponent control of behavior by dorsomedial striatal pathways depends on task demands and internal state. Nat. Neurosci. 25, 345–357. doi: 10.1038/s41593-022-01021-9.

24. Ashwood, Z.C., Roy, N.A., Stone, I.R., Urai, A.E., Churchland, A.K., Pouget, A., and Pillow, J.W. (2022). Mice alternate between discrete strategies during perceptual decision-making. Nat. Neurosci. 25, 201–212. doi: 10.1038/s41593-021-01007-z.

25. Brunton, B.W., Botvinick, M.M., and Brody, C.D. (2013). Rats and humans can optimally accumulate evidence for decision-making. Science 340, 95–98. doi: 10.1126/science.1233912.

26. Kane, G.A., Senne, R.A., and Scott, B.B. (2024). Rat movements reflect internal decision dynamics in an evidence accumulation task. J. Neurophysiol. 132, 1608–1620. doi: 10.1152/jn.00181.2024.

27. Myers, C.E., Interian, A., and Moustafa, A.A. (2022). A practical introduction to using the drift diffusion model of decision-making in cognitive psychology, neuroscience, and health sciences. Front. Psychol. 13, 1039172. doi: 10.3389/fpsyg.2022.1039172.

28. Pohle, J., Langrock, R., van Beest, F.M., and Schmidt, N.M. (2017). Selecting the number of states in hidden markov models: Pragmatic solutions illustrated using animal movement. J. Agric. Biol. Environ. Stat. 22, 270–293. doi: 10.1007/s13253-017-0283-8.

29. Fard, P.R., Park, H., Warkentin, A., Kiebel, S.J., and Bitzer, S. (2017). A bayesian reformulation of the extended drift-diffusion model in perceptual decision making. Front. Comput. Neurosci. 11, 29. doi: 10.3389/fncom.2017.00029.

30. Gupta, D., DePasquale, B., Kopec, C.D., and Brody, C.D. (2024). Trial-history biases in evidence accumulation can give rise to apparent lapses in decision-making. Nat. Commun. 15, 662. doi: 10.1038/s41467-024-44880-5.

31. Powers, M.K., and Green, D.G. (1978). Single retinal ganglion cell responses in the dark-reared rat: grating acuity, contrast sensitivity, and defocusing. Vision Res. 18, 1533–1539. doi: 10.1016/0042-6989(78)90008-1.

32. Johnson, N.P., Gregorich, S.M., and Passaglia, C.L. (2020). Spatiotemporal contrast sensitivity of brown-norway rats under scotopic and photopic illumination. Neuroscience 449, 63–73. doi: 10.1016/j.neuroscience.2020.09.030.

33. Nikbakht, N., and Diamond, M.E. (2021). Conserved visual capacity of rats under red light. Elife 10. doi: 10.7554/eLife.66429.

34. Galindo-Romero, C., Norte-Muñoz, M., Gallego-Ortega, A., Rodríguez-Ramírez, K.T., Lucas-Ruiz, F., González-Riquelme, M.J., Vidal-Sanz, M., and Agudo-Barriuso, M. (2022). The retina of the lab rat: focus on retinal ganglion cells and photoreceptors. Front. Neuroanat. 16, 994890. doi: 10.3389/fnana.2022.994890.

35. Stephenson, R., Lim, J., Famina, S., Caron, A.M., and Dowse, H.B. (2012). Sleep-wake behavior in the rat: ultradian rhythms in a light-dark cycle and continuous bright light: Ultradian rhythms in a light-dark cycle and continuous bright light. J. Biol. Rhythms 27, 490–501. doi: 10.1177/0748730412461247.

36. Yamashita, K., Kinoshita, F.L., Yoshida, S.Y., Matsumoto, K., Mitani, T.T., Fujishima, H., Minami, Y., Morii, E., Yamada, R.G., Okada, S., and Ueda, H.R. (2026). A whole-brain single-cell atlas of circadian neural activity in mice. Science 391, eaea3381. URL: https://doi.org/10.1126/science.aea3381. doi: 10.1126/science.aea3381.

37. Storch, K.F., and Weitz, C.J. (2009). Daily rhythms of food-anticipatory behavioral activity do not require the known circadian clock. Proc. Natl. Acad. Sci. U. S. A. 106, 6808–6813. doi: 10.1073/pnas.0902063106.

38. Ratcliff, R., and Tuerlinckx, F. (2002). Estimating parameters of the diffusion model: approaches to dealing with contaminant reaction times and parameter variability. Psychon. Bull. Rev. 9, 438–481. doi: 10.3758/BF03196302.

39. Vasilev, D., Havel, D., Liebscher, S., Slesiona-Kuenzel, S., Logothetis, N.K., Schenke-Layland, K., and Totah, N.K. (2021). Three water restriction schedules used in rodent behavioral tasks transiently impair growth and differentially evoke a stress hormone response without causing dehydration. eNeuro 8, ENEURO.0424–21.2021. doi: 10.1523/ENEUR0.0424-21.2021.

40. Wood, S.N. (2011). Fast stable restricted maximum likelihood and marginal likelihood estimation of semiparametric generalized linear models: Estimation of semiparametric generalized linear models. J. R. Stat. Soc. Series B Stat. Methodol. 73, 3–36. doi: 10.1111/j.1467-9868.2010.00749.x.

41. R Core Team. R: A Language and Environment for Statistical Computing (2018). URL: https://www.R-project.org/.

42. Navarro, D.J., and Fuss, I.G. (2009). Fast and accurate calculations for first-passage times in wiener diffusion models. J. Math. Psychol. 53, 222–230. doi: 10.1016/j.jmp.2009.02.003.

43. Heathcote, A., Brown, S., and Mewhort, D.J.K. (2002). Quantile maximum likelihood estimation of response time distributions. Psychonomic Bulletin & Review 9, 394–401. doi: 10.3758/BF03196299.

44. Revels, J., Lubin, M., and Papamarkou, T. (2016). Forward-mode automatic differentiation in julia. arXiv [cs.MS]. arXiv:1607.07892.

45. K Mogensen, P., and N Riseth, A. (2018). Optim: A mathematical optimization package for julia. J. Open Source Softw. 3, 615. doi: 10.21105/joss.00615.

46. Bezanson, J., Edelman, A., Karpinski, S., and Shah, V.B. (2017). Julia: A fresh approach to numerical computing. SIAM Rev. Soc. Ind. Appl. Math. 59, 65–98. doi: 10.1137/141000671.

47. Dalle, G. (2024). HiddenMarkovModels.jl: generic, fast and reliable state space modeling. J. Open Source Softw. 9, 6436. doi: 10.21105/joss.06436.

48. Senne, R., Loschinskey, Z., Fourie, J., Loughridge, C., and DePasquale, B.D. (2025). StateSpaceDynamics.jl: A julia package for probabilistic state space models (SSMs). J. Open Source Softw. 10, 8077. doi: 10.21105/joss.08077.

49. Fjelde, T.E., Xu, K., Widmann, D., Tarek, M., Pfiffer, C., Trapp, M., Axen, S.D., Sun, X., Hauru, M., Yong, P., Tebbutt, W., Ghahramani, Z., and Ge, H. (2025). Turing.jl: A general-purpose probabilistic programming language. ACM Trans. Probab. Mach. Learn. 1, 1–48. doi: 10.1145/3711897.

50. Christ, S., Schwabeneder, D., Rackauckas, C., Borregaard, M.K., and Breloff, T. (2023). Plots.jl: A user extendable plotting API for the julia programming language. J. Open Res. Softw. 11. doi: 10.5334/jors.431.

51. Danisch, S., and Krumbiegel, J. (2021). Makie.jl: Flexible high-performance data visualization for julia. J. Open Source Softw. 6, 3349. doi: 10.21105/joss.03349.

52. Wickham, H. (2016). ggplot2: Elegant graphics for data analysis. Use R! 2 ed. Springer International Publishing. URL: https://ggplot2-book.org. doi: 10.1007/978-3-319-24277-4.

53. van der Plas, F., Dral, M., Berg, P., Georgakopoulos, P., Huijzer, R., Bocheński, M., Mengali, A., Burns, C., Lungwitz, B., Priyashan, H., Ling, J., Wu, G., Kadowaki, S., Guinard, C., Zhang, E., Vargas, S.A., Luo, X.R., Schneider, F.S.S., Greimel, F. et al. (2026). JuliaPluto/Pluto.jl: v0.20.25. Zenodo. URL: https://doi.org/10.5281/zenodo.20084276. doi: 10.5281/zenodo.20084276.

